# Discovery and characterization of the first known biological lanthanide chelator

**DOI:** 10.1101/2022.01.19.476857

**Authors:** Alexa M. Zytnick, Sophie M. Gutenthaler-Tietze, Allegra T. Aron, Zachary L. Reitz, Manh Tri Phi, Nathan M. Good, Daniel Petras, Lena J. Daumann, N. Cecilia Martinez-Gomez

## Abstract

Many bacteria secrete metallophores, low-molecular weight organic compounds that bind ions with high selectivity and affinity, in order to access essential metals from the environment.^1^ Previous work has elucidated the structures and biosynthetic machinery of metallophores specific for iron, zinc, nickel, molybdenum, and copper.^1^ No lanthanide-specific metallophore has been discovered despite the knowledge that lanthanide metals (Ln) have been revealed to be essential cofactors for certain alcohol dehydrogenases across a diverse range of phyla.^2^ Here, we report the biosynthetic machinery, the structure, and the physiological relevance of the first known lanthanophore, methylolanthanin. The structure of methylolanthanin exhibits a unique 4-hydroxybenzoate moiety which has not previously been described in other metallophores. We find that production of methylolanthanin is required for normal levels of Ln accumulation in the methylotrophic bacterium *Methylobacterium extorquens* AM1, while overexpression of the molecule greatly increases bioaccumulation. Our results provide a clearer understanding of how Ln-utilizing bacteria sense, scavenge, and store Ln; essential processes in the environment where Ln are poorly bioavailable. Beyond Ln, we anticipate our study to be a starting point for understanding how organisms acquire other *f*-block metals, the actinides.^3^ More broadly, the discovery of a lanthanophore opens doors for study of how biosynthetic gene clusters are repurposed for new functions, how metallophores acquire their metal specificity, and the complex relationship between metal homeostasis and fitness.

## MAIN

Metal ions are essential for life—it is estimated that 40-50% of all enzymes require a metal ion for proper function ^4–6^. Whether serving in catalytic or structural roles, metal ions are involved in biological processes that range from respiration and DNA replication to the biosynthesis of metabolic intermediates. The importance of metal ions is underscored by the extensive systems that organisms across the entire tree of life have developed to not only sense and acquire metals from the environment but also regulate concentrations within their cells.

Many bacteria secrete metallophores, small-molecule chelators, to make environmental metals more bioavailable (4). Siderophores (Fe), chalkophores (Cu), zincophores (Zn), molybdophores (Mo), and nickelophores (Ni) are metallophores that have been reported.^1^ Metallophore structures are categorized based on their metal-binding moieties—such as catecholates, phenolates, hydroxamates, carboxylates, and diazeniumdiolates—each of which have unique physiochemical properties.^7,8^ The majority of known metallophores are biosynthesized by nonribosomal peptide synthetases (NRPSs) or NRPS-independent siderophore (NIS) synthetases. Metallophore biosynthetic gene clusters (BGCs) often include genes encoding transport systems that bring the metallophore-metal complex into the cell for metal release. In Gram-negative bacteria, TonB complexes provide the energy for transport into the periplasm *via* beta-barrel outer membrane receptors. Depending on the metallophore system, the complex may remain in the periplasm or be transported to the cytoplasm *via* an ATP-binding cassette (ABC) transporter before the metal is released by metallophore hydrolysis and/or metal reduction. As free metal in the cell can be toxic due to the formation of reactive oxygen species or through mismetallation of enzymes, metal acquisition and uptake genes are often tightly regulated by metal-responsive transcription factors such as ferric uptake regulator (Fur) and nickel uptake regulator (Nur) family proteins or by sigma factor / anti-sigma factor signaling systems.^9–11^

While many are familiar with canonical *d*-block life metals such as iron, copper, and magnesium, members of the *f-*block lanthanide series have recently solidified their place among the life metals. Lanthanides were found to be biologically relevant in 2011 when it was discovered that MxaFI, a pyrroloquinoline quinone (PQQ) and Ca-dependent methanol dehydrogenase from the methylotrophic bacterium *Methylobacterium extorquens* AM1, had a Ln-dependent homolog, XoxF.^12^ While MxaFI and XoxF have similar active sites, crystal structures confirmed that the XoxF active site contains an extra aspartic acid residue that is essential for Ln binding.^13^ Further studies found that Ln-dependent enzymes are not isolated to methylotrophic bacteria nor to exhibiting methanol dehydrogenase activity; the ethanol dehydrogenase ExaF from *M. extorquens* AM1 as well as the ethanol dehydrogenase PedH from *P. putida* KT2440 were recently discovered to be Ln-dependent.^14,15^

All known Ln-dependent enzymes to date are periplasmic proteins, but the details of Ln acquisition, transport, and integration into enzymes are not fully understood. The Ln utilization and transport (*lut*) cluster that was discovered in *M. extorquens* AM1^16^ and the closely related *M. extorquens* PA1 is required for Ln-dependent growth.^17^ In this system Ln are initially brought into the periplasm through the TonB-dependent receptor LutH. There is currently no experimental evidence of free Ln in the periplasm. Rather, Ln are then transported to the cytoplasm via the *lut*-encoded ABC transport system, but the necessity of Ln reaching the cytoplasm is not yet understood. While it seems that the *lut* cluster encodes the primary system for Ln uptake, it remains unknown whether other transport systems are also used.

The presence of a Ln-chelating metallophore has long been proposed.^17,18^ Methylotrophs like *M. extorquens* AM1 are known to thrive in mesophilic environments that are rich in poorly soluble Ln such as the soil and the phyllosphere (the arial region of plants) where a mechanism for solubilization would be necessary for Ln-dependent methylotrophic growth. Concentrations of Ln in the phyllosphere range from 0.7 μg/g dry weight to 7 μg/g dry weight, but bioavailable Ln concentrations are likely far less.^17,19^ In most soil environments Ln are found in highly insoluble oxide and phosphate forms, usually in minerals and ores such as monazite and bastnäsite.^20^ Despite this, most work done in a laboratory setting has used soluble chloride salts to understand Ln biology.

In this study, we utilize Ln sources of high (NdCl_3_) and low (Nd_2_O_3_) solubility to identify and structurally characterize the first known metallophore used to obtain Ln, a lanthanophore we have named methylolanthanin (MLL, **1**). We identified the *mll* biosynthetic gene cluster through assessment of *M. extorquens* AM1’s transcriptional response to a low solubility Ln source. We find overexpression of the MLL biosynthetic genes *in trans* increases growth and Ln bioaccumulation, while deletion of these genes generates a severe defect in Ln bioaccumulation. Furthermore, we demonstrate the Ln-binding of MLL and find that exogenous addition of the compound to a mutant lacking the capability to biosynthesize it improves growth yield.

## Unbiased discovery of Ln solubility gene networks

To better understand the impacts of Ln solubility on methylotrophy in *M. extorquens* AM1, we measured methanol growth with soluble NdCl_3_ and poorly soluble Nd_2_O_3_ using Δ*mxaF*, a strain that lacks catalytically active MxaFI and is therefore dependent on Ln for growth on methanol. Higher Ln solubility resulted in significantly faster growth (*p* value = 0.001) (Fig. 1a). We employed RNA-seq transcriptomics to gain unbiased insight into the reasons for this significant growth defect with Nd_2_O_3_. We identified 1,468 differentially expressed genes between the NdCl_3_ and Nd_2_O_3_ conditions (Fig. 1b).

**Fig. 1.**
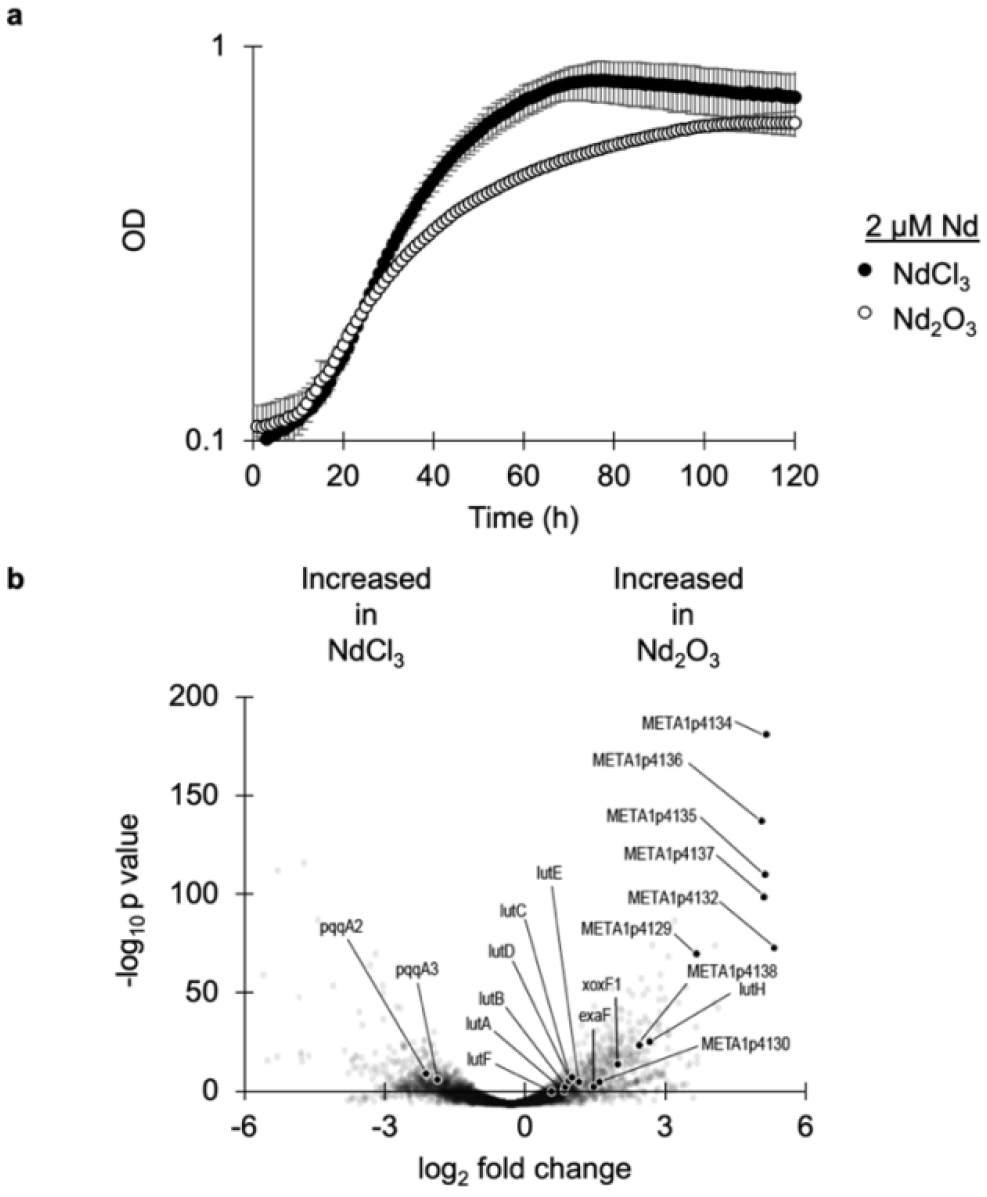
Differences in Ln source solubility affect growth and transcriptional response of *M. extorquens*. **a**, Growth rate of Δ*mxaF* is significantly increased during growth on soluble NdCl_3_ (filled circles) compared to poorly soluble Nd_2_O_3_ (open circles). Individual data points represent the mean of three replicates. **b**, Volcano plot of DEGs comparing NdCl_3_ and Nd_2_O_3_ conditions with biosynthetic genes for methylolanthanin highlighted in gray. Individual data points represent the mean of three replicates.

When Nd_2_O_3_ was provided, the gene encoding the primary Ln-dependent MDH (*xoxF1*) was among the most upregulated genes, exhibiting 5-fold upregulation when compared to the soluble Ln condition (Fig. 1b). This result corroborates previous findings on the upregulation of *xox* genes generated by the presence of exogenous LaCl_3_ ^21–24^. We also found the gene encoding the Ln-dependent ethanol dehydrogenase *exaF* was shown to have a 3-fold increase in expression during growth with Nd_2_O_3_ compared to growth with NdCl_3_, suggesting that alternative Ln alcohol dehydrogenases are important when Ln are less bioavailable.

Previous studies have shown that although upregulation of *xoxF1* occurs as a response to soluble Ln, concomitant upregulation of the PQQ biosynthetic genes does not occur ^21,23^. We find similar results in this study, as most of the genes encoding the PQQ biosynthetic machinery were not significantly upregulated in the NdCl_3_ condition. Interestingly, the orphan *pqqA* copies *pqqA2* and *pqqA3* were upregulated 4-fold in the NdCl_3_ condition compared to the Nd_2_O_3_ condition (Fig. 1b).

The *lut* cluster, which encodes the TonB dependent transporter LutH, an ABC transport system, and various periplasmic proteins, is essential for light Ln transport. Previous work has shown expression of *lutH* does not change in response to soluble LaCl_3_ ^21^.In this study, *lutH* was downregulated 9-fold in the NdCl_3_ condition compared to the no Ln condition and upregulated 9-fold compared to the Nd_2_O_3_ condition, denoting tight regulation of the TonB dependent transporter based on Ln species (Fig. 1b). The remaining *lut* genes exhibited 2-fold upregulation on average during growth with Nd_2_O_3_.

### Identification of a putative lanthanophore locus

Of all genes identified in RNAseq analysis, the genes META1p4129 through META1p4138 were the most highly upregulated in the Nd_2_O_3_ condition, with an average increase in expression of 32-fold compared to growth with NdCl_3_ (Fig. 1b, Table 1). The locus, which we have named *mll* (for methylolanthanin, *vide infra*), is homologous to BGCs responsible for the transport, regulation, and biosynthesis of NRPS-independent siderophores containing citrate and 3,4-dihydroxybenzoate (3,4-DHB) chelating groups, including rhodopetrobactin, petrobactin, and roseobactin (Fig. 2, Extended Data Fig. 1).^25^ META1p4132-4135 (*mllA, mllBC, mllDE*, and *mllF*) are homologous to the well-studied petrobactin locus in *B. subtilis, asbABCDEF*,^26^ differing by two sets of gene fusions; the fusion between *asbD* and *asbE* homologs is also present in the rhodopetrobactin and roseobactin loci, while the *mllBC* fusion has not been observed in characterized homologs. The presence of putative acetyltransferase *mllH* (META1p4137) suggested the incorporation of an acetylated (homo)spermidine linker, as seen in rhodopetrobactin. META1p4129-4131 (*mluARI*, **m**ethylo**l**anthanin **u**ptake) putatively encode, respectively, a TonB-dependent outer membrane receptor, an anti-sigma factor, and a sigma factor. Homologous proteins form a cell-surface signaling pathway, where the import of a ferric siderophore induces further expression of the receptor.^27^ Two domains of unknown function are encoded in the cluster. META1p4136 (*mllG*) belongs to DUF2218 and is also present in the rhodopetrobactin locus. Homologous VCA0233 from *Vibrio cholerae* neighbors an iron uptake regulator and xeno-siderophore uptake genes,^28^ suggesting DUF2218 is involved in regulation or transport rather than biosynthesis. META1p4138 (*mllJ*) belongs to ferritin-like DUF4142 and is putatively exported into the periplasm. A phylogenetic tree of representative *Methylorubrum* and *Methylobacterium* genomes was reconstructed using PhyloPhlAn, and ITOL was used to map the presence of homologous BGCs, as determined by a custom version of antiSMASH.^29–31^ The majority of *Methylorubrum* species, forming a single clade, contain homologs of the *mll* locus, as well as *Methylobacterium currus* TP3 and *Methylobacterium aquaticum* BG2 (Extended Data Fig. 1). Together, the insights that the *mll* locus is (1) upregulated during growth with an insoluble Ln source, (2) contains genes commonly found in metallophore BGCs, and (3) is conserved across many *Methylorubrum* strains, all suggested that the cluster produces a rhodopetrobactin-like lanthanophore.

**Table 1.** Comparison of methylolanthanin and rhodopetrobactin BGCs.

| Gene | Locus tag | log2 fold change | q-value | Putative product | Rhodopetrobactin BGC homolog | % ID |
| --- | --- | --- | --- | --- | --- | --- |
| *mluA* | META1p4129 | 4.2 | 3.70E-84 | TonB dependent receptor | Rpal_2620 | 62 |
| *mluR* | META1p4130 | 2 | 3.00E-12 | Anti-sigma factor | Rpal_2623 | 30 |
| *mluI* | META1p4131 | 6.1 | 3.60E-58 | Sigma factor | Rpal_2633 | 30 |
| *mllA* | META1p4132 | 6 | 1.20E-87 | *asbA*-like NIS synthetase | Rpal_2632 | 50 |
| *mllBC* | META1p4133 | 6.3 | 1.10E-165 | *asbB*-like NIS synthetase and | Rpal_2631 | 52 |
|  |  |  |  | *asbC*-like adenylation domain | Rpal_2630 | 56 |
| *mllDE* | META1p4134 | 5.8 | 3.60E-209 | *asbD*-like aryl carrier protein and<br>*asbE*-like ligase fusion | Rpal_2629 | 54 |
| *mllF* | META1p4135 | 5.8 | 8.30E-130 | *asbF*-like 3,4-DHB synthase | Rpal_2628 | 52 |
| *mllG* | META1p4136 | 5.7 | 3.20E-160 | DUF2218 family protein | Rpal_2619 | 67 |
| *mllH* | META1p4137 | 5.8 | 6.80E-117 | GNAT family acetyltransferase | Rpal_2621 | 74 |
| *mllJ* | META1p4138 | 2.9 | 6.10E-33 | DUF4142 family protein | - | - |

**Fig. 2.**
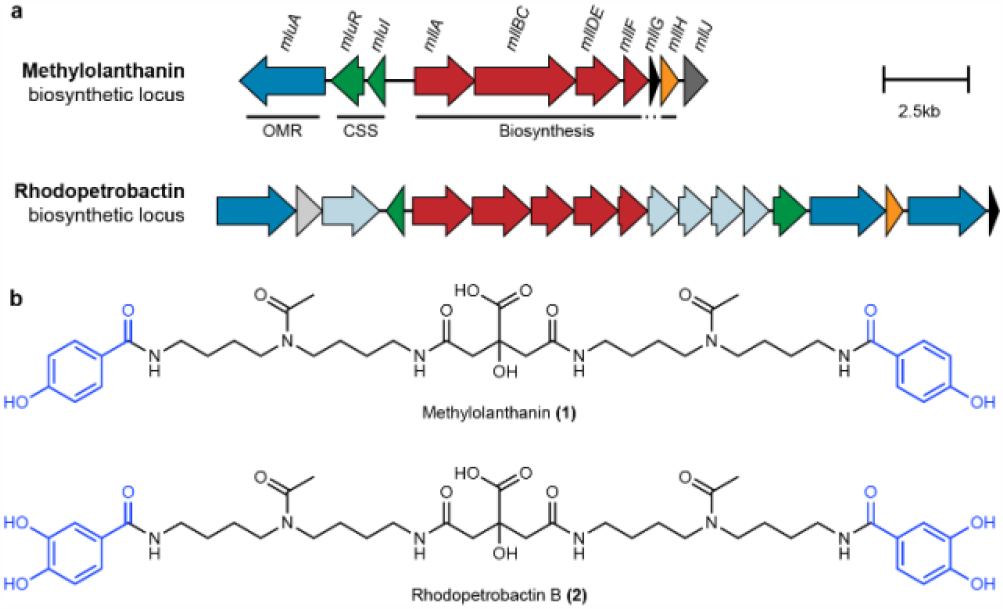
Biosynthetic gene clusters and chemical structures of methylolanthanin (1) and the related siderophore rhodopetrobactin (2). **a**, The methylolanthanin BGC from *M. extorquens* AM1 (*mll/mlu*, META1p4129-4138) and the rhodopetrobactin BGC from *R. palustris* TIE-1.^25^ Homologous pathways between BGCs share the same color. Genes are drawn to scale. OMR, TonB-dependent outer membrane receptor; CSS, cell surface signaling. Additional homologous clusters are presented in Extended Data Fig. 1. **b**, Chemical structures of methylolanthanin (1) and rhodopetrobactin (2), which differ by the presence of 4-HB versus 3,4-DHB, respectively (in blue).

### Structural and functional elucidation of methylolanthanin

To determine the identity of the putative lanthanophore produced by the *mll* cluster, mutants lacking (Δ*mxaF*Δ*mll*) and overexpressing (Δ*mxaF*/pAZ1) META1p4132 through META1p4138 were constructed. The genome of the *mll* deletion strain was sequenced, and it was confirmed that no additional mutations had been acquired. Supernatant extracts were analyzed using ultra-high performance liquid chromatography tandem mass spectrometry (UPLC-MS/MS) in both positive and negative ionization modes. Classical molecular networking using the Global Natural Products Social (GNPS) Molecular Networking platform^32^ revealed only one molecular family that was found in Δ*mxaF* supernatants and not in Δ*mxaF*Δ*mll* supernatants, containing a compound with mass to charge ratio (*m/z*) of 799.4232 in positive mode and 797.4092 in negative mode (Fig. 3a and Extended Data Fig. 2). Multivariate and univariate (Fig. 3b,c and Extended Data Fig. 2) statistical analyses were performed in parallel on blank subtracted, normalized, and imputed feature tables processed after feature finding in MZmine 3.^33^ Volcano plot analysis comparing *ΔmxaF* supernatant with Δ*mxaF*Δ*mll* supernatants was consistent with classical molecular networking results, revealing that the putative *mll* product (positive mode *m/z* 799.4232; negative mode *m/z* 797.4092) illustrates the second highest fold-change and most significant p-value in negative mode data (Fig. 3d), and the putative *mll* product is differentially abundant across wild type, knockout and overexpression samples.

**Fig. 3.**
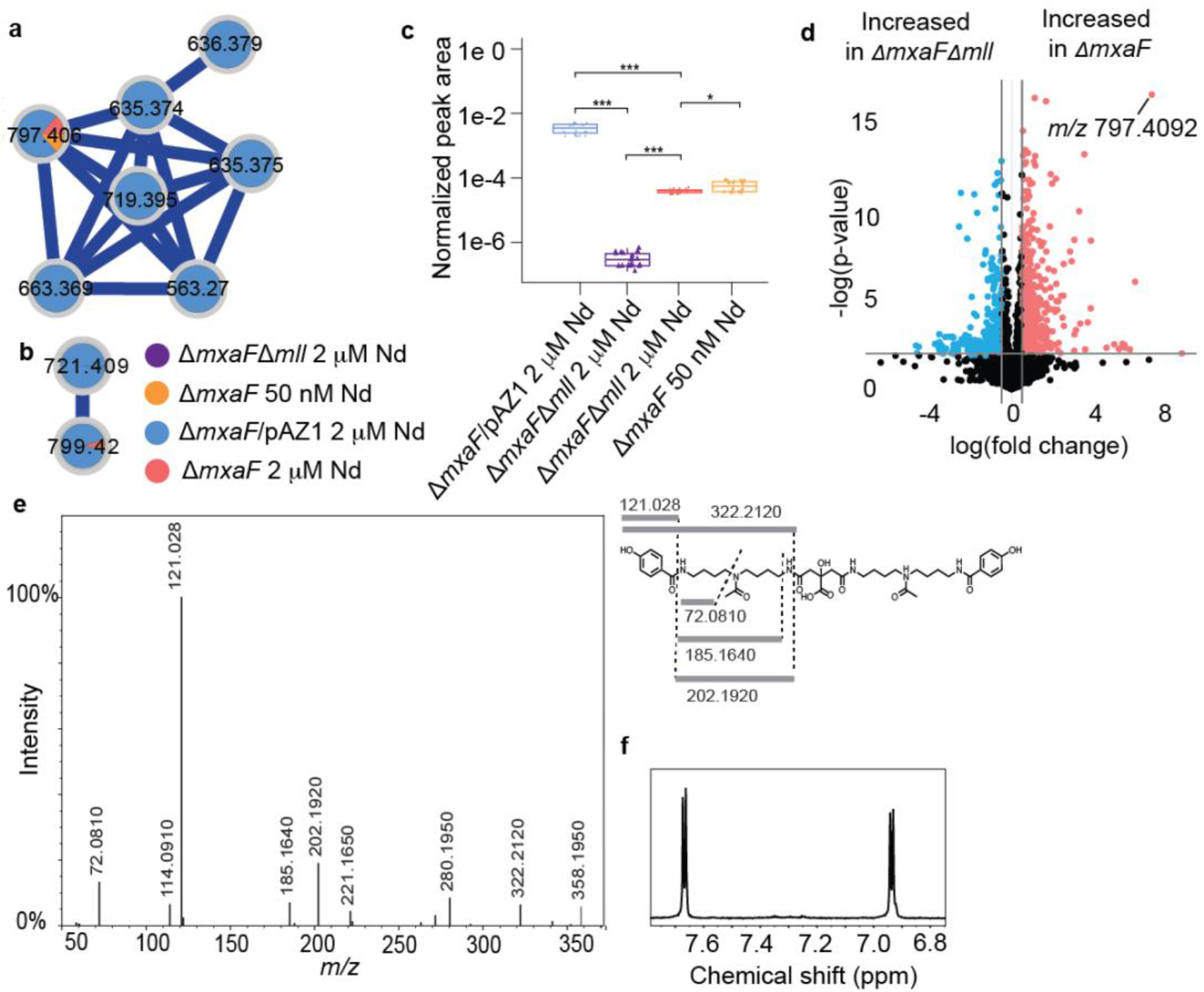
Identification of methylolanthanin via molecular networking and MS/MS. **a**, ESI-UPLC-MS/MS in negative ionization mode reveals a molecular family of MS/MS spectra only found in Δ*mxaF* (orange and red) and Δ*mxaF*/pAZ1 (blue) but not in Δ*mxaF*Δ*mll* (purple). **b**, ESI-UPLC-MS/MS in positive ionization mode reveals a molecular family of MS/MS spectra only found in *ΔmxaF* (orange and red) and *ΔmxaF*/pAZ1 (blue) but not in Δ*mxaF*Δ*mll* (purple). Two features (*m/z* 799.4232 in positive mode / *m/z* 797.4092 in negative mode and *m/z* 721.4114 in positive mode / *m/z* 719.3978 in negative mode) are found in both positive and negative ionization mode molecular families. **c**, Boxplots of the normalized peak area of feature *m/z* 797.4092 in negative mode reveal significant differences between peak area in all conditions. Statistics were calculated with n=20 per group Kruskal-Wallis, followed by pairwise Wilcoxon tests and Benjamini-Hochberg (BH) correction (**p< 0.001, *p< 0.05). **d**, *m/z* 797.4092 is one of the most significantly increased features in Δ*mxaF* versus Δ*mxaF*Δ*mll* supernatants when analyzed using ESI-UPLC-MS/MS in positive ionization mode. Volcano plot analysis was performed with n=5 per group. Fold changes 0.5 and a p-value < 0.01 are highlighted. **e**, The consensus MS/MS spectrum in positive mode is consistent with the proposed structure, as highlighted by the key fragments in the chemical structure. **f**, The NMR aromatic region of isolated methylolanthanin supports a para-substituted mono-hydroxybenzoate moiety (^1^H NMR spectrum, 800 MHz, D_2_O + 0.003% TMSP).

Tandem mass spectrometry (MS/MS) data collected in both negative and positive modes revealed fragments that fit with a proposed structure that resembles rhodopetrobactin B minus two oxygen atoms. Key fragments in positive mode consensus MS/MS include *m/z* 322.2120, 202.1920, 185.1640, 121.0280, and 72.0810 (Fig. 3e). These fragments indicate the (Fig. 3e). These fragments indicate the presence of acetylated homospermidine (4,4’-diaminodibutylamine) and monosubstituted hydroxybenzoic acid (HB). The presence of two HB moieties account for the mass difference with rhodopetrobactin B, which instead contains two 3,4-dihydroxybenzoic acid (3,4-DHB) moieties. The neutral loss of 156.006 (between fragments at *m/z* 299.1590 and 143.1540) is consistent with the presence of citrate.

The molecule was isolated to confirm the proposed structure and determine the position of the hydroxyl group in the hydroxybenzoic acid. NMR spectroscopy of the purified compound revealed two doublets in the aromatic region, clearly showing the para substitution of the hydroxybenzoic acid moiety (Fig. 3f). Thus, the structure of the molecule, which we named methylolanthanin (MLL, **1**), was fully established and confirmed by NMR (Extended Data Fig. 3): a central citrate group is linked to two 4-HB moieties *via* homospermidine residues, each of which is acetylated at the central amine (Fig. 2b). Signals in the ^1^H NMR spectrum show varying degrees of unexpected splitting that can be explained by the presence of four conformers of roughly equal abundance stemming from different orientations of the two HB moieties account for the mass difference with rhodopetrobactin B, which instead contains two 3,4-dihydroxybenzoic acid (3,4-DHB) moieties. The neutral loss of 156.006 (between fragments at *m/z* 299.1590 and 143.1540) is consistent with the presence of citrate.

The molecule was isolated to confirm the proposed structure and determine the position of the hydroxyl group in the hydroxybenzoic acid. NMR spectroscopy of the purified compound revealed two doublets in the aromatic region, clearly showing the para substitution of the hydroxybenzoic acid moiety (Fig. 3f). Thus, the structure of the molecule, which we named methylolanthanin (MLL, **1**), was fully established and confirmed by NMR (Extended Data Fig. 3): a central citrate group is linked to two 4-HB moieties *via* homospermidine residues, each of which is acetylated at the central amine (Fig. 2b). Signals in the ^1^H NMR spectrum show varying degrees of unexpected splitting that can be explained by the presence of four conformers of roughly equal abundance stemming from different orientations of the two acetyl groups (Extended Data Fig. 4). The diastereotopic methylene groups in the citrate moiety show ^1^H NMR signals for four overlapping AB systems instead of two doublets with a roof effect (Extended Data Figs. 3 and 4). The overlap of the signals depends on the solvent system used (H2O/ D2O vs. D2O; Extended Data Fig. 4). The signals of the secondary amine protons (only observed in H2O/D2O) also showed additional splitting most probably caused by the four possible conformers (see Extended Data Fig 4 for deconvoluted signals).

Furthermore, we were able to show *via* direct injection mass spectrometry that methylolanthanin is capable of binding lanthanides of varying sizes by co-injecting La(III), Nd(III), and Lu(III). In each case, a a complex of the formula [MLL-H^+^+Ln^3+^]^2+^ can be found with the corresponding isotopic pattern of La, Nd and Lu (Fig. 4).

**Fig. 4.**
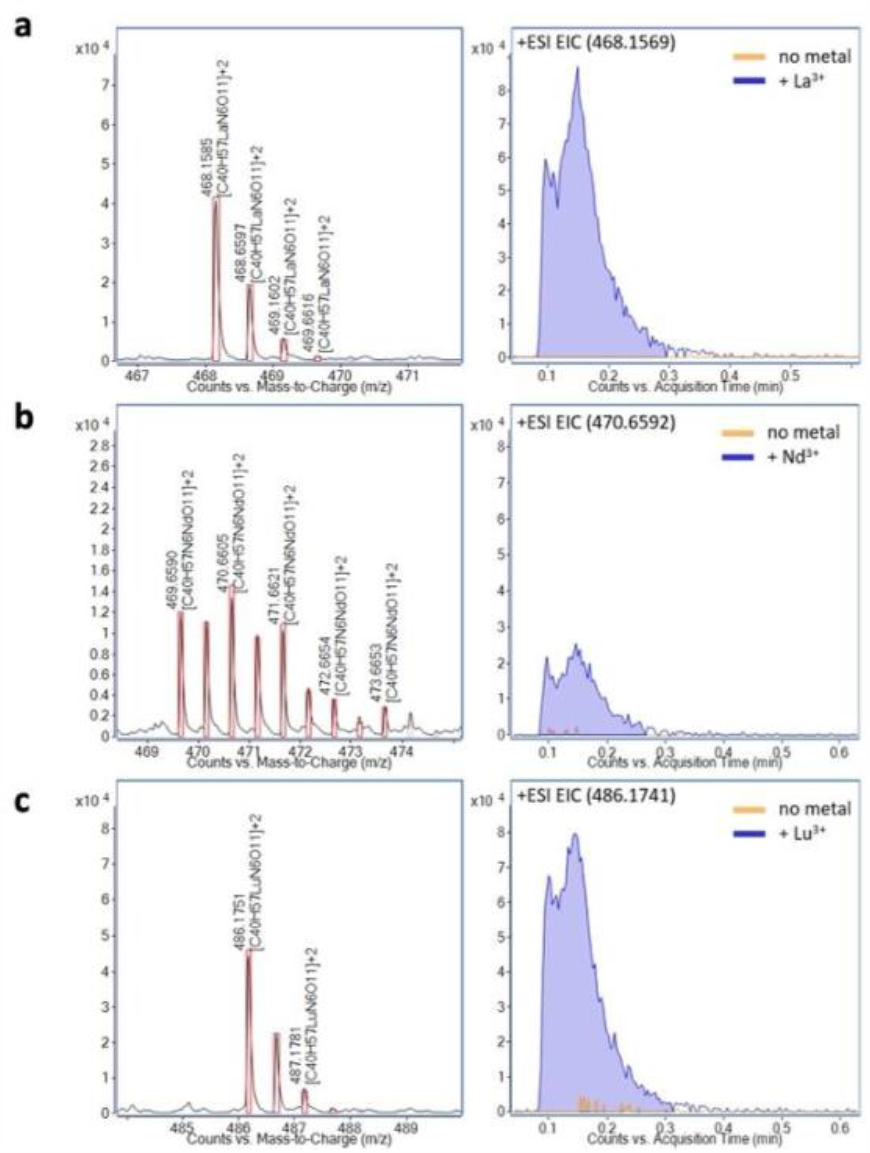
Methylolanthanin binds lanthanum (La), neodymium (Nd), and lutetium (Lu) EICs of methylolanthanin-Ln^3+^ complexes and corresponding mass spectra showing the isotopic pattern supporting complex formation. **a**, La(III). **b**, Nd(III). **c**, Lu(III). Theoretical isotopic patterns are indicated by boxes.

### Phenotypic analysis of methylolanthanin mutants

To better understand the effect methylolanthanin has on the physiology of *M. extorquens* AM1, the growth and Ln bioaccumulation of Δ*mxaF*Δ*mll* and Δ*mxaF*/pAZ1 were studied alongside a Δ*mxaF* control. As shown above, Δ*mxaF* exhibited a clear defect when grown with Nd_2_O_3_ compared to when grown with NdCl_3_ (Fig. 1a). This defect was rescued when the *mll* cluster was overexpressed; Δ*mxaF*/pAZ1 Nd_2_O_3_ cultures grew at a rate of 0.026 hr^-1^, a nearly 50% increase compared to the growth of Δ*mxaF* with Nd_2_O_3_ and a growth rate closer to that of Δ*mxaF* on NdCl_3_ (0.037 hr^-1^) (Fig. 5a). Interestingly, Δ*mxaF*/pAZ1 cultures incurred a defect in growth rate when compared to Δ*mxaF* when grown with soluble NdCl_3_ (Fig. 5b). Pure methylolanthanin (50 nM), isolated from Δ*mxaF*/pAZ1 cultures, was added to Δ*mxaF* or Δ*mxaF*Δ*mll* growing with 2 μM NdCl_3_ to assess the effect of the molecule on growth. Presence of exogenous methylolanthanin significantly increased growth yield (maximal OD in stationary phase) of both cultures (*p* values = 0.036 and 0.037) (Fig. 5c).

**Fig. 5.**
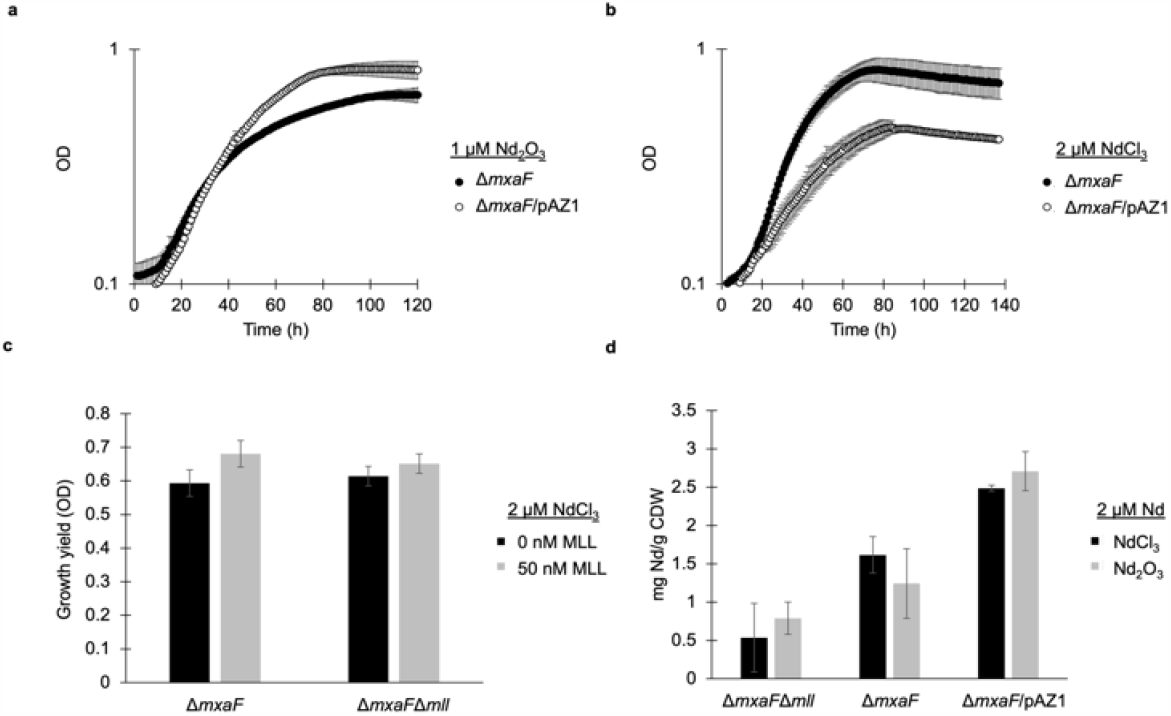
Manipulation of *mll* affects growth yield and neodymium bioaccumulation. **a**, Growth of Δ*mxaF* (filled circles) and Δ*mxaF*/pAZ1 (empty circles) with 1 μM Nd_2_O_3_. Individual data points represent the mean of three replicates. **b**, Growth of Δ*mxaF* (filled circles) and Δ*mxaF*/pAZ1 (empty circles) with 2 μM NdCl_3_. Individual data points represent the mean of three replicates. **c**, Growth yield of Δ*mxaF* and Δ*mxaF*Δ*mll* growing with 2 μM NdCl_3_ with or without 50 nM MLL. Values represent the mean of three replicates. **d**, Intracellular accumulation of Nd by Δ*mxaF*Δ*mll*, Δ*mxaF*, and Δ*mxaF*/pAZ1 with 2 μM NdCl_3_ and 1 μM Nd_2_O_3_. Values represent the mean of three replicates.

As Ln-dependent growth of *M. extorquens* AM1 has previously been shown to be tied to Ln bioaccumulation, inductively coupled plasma mass spectrometry (ICP-MS) was used to measure intracellular concentrations of Nd within Δ*mxaF*, Δ*mxaF*Δ*mll*, and Δ*mxaF*/pAZ1 grown with Nd_2_O_3_ or NdCl_3_.^16,34^ Δ*mxaF* bioaccumulated Nd to levels previously reported for other light Ln (Fig. 5d).^34^ Deletion of *mll* decreased Nd bioaccumulation with both Ln sources, notably by 1.8 fold in the NdCl_3_ condition, while overexpression of *mll* increased Nd bioaccumulation by 3.5 fold on average (Fig. 5d).

## DISCUSSION

Metallophores for iron, zinc, copper, nickel, and molybdenum have been chemically and biosynthetically elucidated. No Ln-specific metallophore (lanthanophore) has been discovered even as Lns are essential cofactors for certain alcohol dehydrogenases.^2^ Here we report the chemical structure, biosynthesis and physiological relevance for methylolanthanin (**1**), a first-of-its-class Ln-binding metallophore. The narrow taxonomic distribution of *mll* (Extended Data Fig. 1) and the common function of homologous molecules as Fe-binding siderophores suggest that the methylolanthanin pathway entered *Methylorubrum* through horizontal gene transfer and evolved to chelate Ln in place of Fe, although further study is required to confirm this hypothesis.

The production of metallophores benefits the organisms that produce them and the “cheaters’’ that utilize them.^35,36^ We can infer that methylolanthanin provides a benefit for producers in environments where Ln are poorly bioavailable, such as the soil and phyllosphere, whereas Ln-utilizing extremophiles in Ln richly bioavailable environments would not require the production of such a molecule. Surveying the RefSeq non-redundant protein record for organisms that encode clusters like the *mll* cluster, we find similar clusters in mesophilic methylotrophic Ln-utilizing organisms but not in extremophilic, methylotrophic Ln-utilizing organisms (*Methylacidiphilum fumariolicum* SolV, isolated from a volcanic mudpot) (Extended Data Fig. 1).

Growth of Δ*mxaF*Δ*mll* (Extended Data Fig. 5) denotes that production of methylolanthanin is not essential under the conditions tested. This leaves open questions about alternative systems for Ln uptake, perhaps through non-specific ion transporters or through the production of other chelation systems. Previous work has shown extracts of a *M. extorquens* strain evolved to use the heavy Ln Gd have exhibited a 6-fold increase in PQQ levels compared to Δ*mxaF* extracts.^34^ Additionally, PQQ has been shown to be able to bind Ln.^37,38^ While in this study we did not observe an increase in the expression of most PQQ biosynthetic genes during growth with poorly bioavailable Nd_2_O_3_, we did observe increased expression of two orphan *pqqA* genes (Fig. 1b). *pqqA*, encoding the peptide precursor to PQQ, has previously been reported to be nonessential for PQQ biosynthesis in *M. extorquens* AM1, but orphan *pqqA* copies were not investigated in these studies.^39^ In *Methylovorus* sp. MP688, which is equipped with five *pqqA* copies, the availability of PqqA has been shown to be a rate determining step in PQQ biosynthesis.^40^ Taken together, it is possible that PqqA could be a component of an alternative Ln uptake system.

It is known that *M. extorquens* AM1 is responsive to Ln levels as low as 2.5 nM, but it is not yet known why Ln are bioaccumulated at much higher levels.^41^ *M. extorquens* AM1 is a candidate to recover lanthanides selectively from electronic waste and old magnets^42^. Previously it has been shown that *M. extorquens* AM1 can selectively bioaccumulate Nd from NdFeB magnets, hence neodymium salts were chosen in this study. Clearly, *M. extorquens* AM1 stores far more Ln than it requires for robust growth. Cultures of Δ*mxaF*Δ*mll* exhibit a 30% decrease in bioaccumulation yet can grow similarly to Δ*mxaF* cultures that exhibit typical levels of Ln bioaccumulation (Fig 5d and Extended Data Fig. 5). Further research is needed to understand the complex relationship between growth and Ln storage.

The most striking feature of methylolanthanin is the 4-HB moiety, which has not been reported in any characterized metallophore. Baars *et al*. reported small amounts of rhodopetrobactin analogs from *R. palustris*, rhodopetrobactins A - O, A - 2O, and B - O, that also contain one or two HB moieties (possibly 4-HB) in place of 3,4-DHB. The molecular weight of methylolanthanin would be equivalent to “rhodopetrobactin B - 2O”, which was not observed by Baars *et al*. Conversely, we did not observe any putative 3,4-DHB–containing analogues in the *M. extorquens* AM1 supernatant. The presence of 4-HB in a metallophore is unexpected, as most metallophore structures contain bidentate chelating moieties. Evolution may have favored this unusual structure to obtain sufficient lanthanides for the needs of the cell while avoiding toxic levels of iron accumulation. We have shown elsewhere that Fe bioaccumulation remains unaffected by overexpression of *mll*.^42^ The Ln affinity of methylolanthanin may be stalled at a local fitness peak, where evolving greater affinity for Ln binding would incur a fitness disadvantage that cannot occur under selection. Siderophores, zincophores, and other metallophores exhibit convergent evolution, where biosynthetically unrelated molecules share the same function. The existence of methylolanthanin suggests that other lanthanophores may have evolved, possibly with higher lanthanide affinity and selectivity.

## METHODS

### General working and culturing conditions

MilliQ grade water and HPLC grade solvents were used in this work, unless otherwise stated. For mass spectrometry and related experiments LCMS grade solvents were used. Culturing occurred under sterile conditions. *M. extorquens* AM1 strains (WT Rif^R^ derivative^43^, Δ*mxaF*^*44*^, Δ*mxaF*/pAZ1, and Δ*mxaF*Δ*mll*) were grown in Hypho medium^45^ (K_2_HPO_4_ 14.5 mM, NaH_2_PO_4_ 18.8 mM, MgSO_4_*7 H_2_O 0.8 mM, (NH_4_)_2_SO_4_ 3.8 mM, FeSO_4_*7 H_2_O 3.69 μM, Na_2_*EDTA*2 H_2_O 26.86 μM, CaCl_2_*2 H_2_O 9.98 μM, MnCl_2_*4 H_2_O 5.11 μM, (NH_4_)_6_Mo_7_O_2_ 0.18 μM, CuSO_4_ 1.26 μM, CoCl_2_*6 H_2_O 6.72 μM, ZnSO_4_*7H_2_O 15.3 μM) but reducing the phosphate concentration by half, from 33.3 mM to 16.6 mM (referred to as ½ Hypho). 3 mL cultures with 15 mM succinate (MilliporeSigma) were grown overnight in 15 mL round-bottom glass culture tubes (Thermo Fisher Scientific) at 30 °C and 200 rpm and then diluted into the desired volume of fresh½ Hypho to an OD600 of 0.01. When necessary, sucrose, kanamycin, chloramphenicol, ampicillin, and methylolanthanin were added to final concentrations of 5%, 50 μg/mL, 12.5 μg/mL, 50 μg/mL, and 50 nM, respectively. Growth curves were obtained with 650 μL cultures grown in transparent 48 well plates (Corning) incubated at 30 °C and 548 rpm using a Synergy HTX plate reader (Biotek). The above conditions were used unless otherwise noted.

### RNA sequencing

50 mL ½ Hypho media supplemented with 50 mM methanol and 1 μM Nd_2_O_3_ (99.99%, Chempur) or 2 μM NdCl_3_ (99.99%, MilliporeSigma) was used to culture WT AM1 and Δ*mxaF* in 250 mL glass Erlenmeyer flasks to mid-exponential phase (OD600 = 0.6). Total RNA samples were generated and quality was corroborated as described previously.^46^ rRNA depletion using the Ribo-Zero RNA Plus rRNA Depletion Kit (Illumina), library preparation, and Illumina Hi-Seq sequencing were performed by the Microbial Genome Sequencing Center (MiGS). Using KBase^47^, reads were aligned with HISTAT2, transcripts were assembled with StringTie, and DEGs were identified using DESeq2.

### Comparative genomics

Gene cluster functions were predicted using hmmscan (EMBL webserver)^48,49^ and through NCBI BLAST^50^ matches against the rhodopetrobactin BGC from *Rhodopseudomonas palustris* TIE-1.^25^ Genomes were retrieved from NCBI RefSeq^51^ in Genbank and protein fasta formats using ncbi-genome-download v0.3.3 for phylogenetic analysis.^52^ The following genomes were selected: complete genomes from *Methylorubrum* and *Methylobacterium*, draft representative genomes from *Methylorubrum*, and three additional *Methylobacteriaceae* genomes as an outgroup (Extended Data Fig. 1, Supplementary Table 1). The phylogenetic tree was reconstructed from conserved protein groups using PhyloPhlAn v3.0.67^29^ with the following configuration parameters: “-d a --msa mafft --tree1 raxml --db_aa diamond --map_aa diamond” and “--diversity medium”.^53–55^ Homologs of the methylolanthanin locus were found using a custom version of antiSMASH v7.0.1^31^ (https://doi.org/10.5281/zenodo.8348225), detection rule for homologs of the petrobactin biosynthetic genes *asbABCDE*. A BGC region met the “AsbABCDE” rule if it contained matches to existing antiSMASH models for IucA_IucC (*asbA/asbB*), AMP-binding (*asbC*), and PP-binding (*asbD*), as well as the Pfam model for DUF6005 (*asbE*, PF19468.2).^56^ No *asbF* model was used due to a low bitscore between *mllF* and the expected Pfam model PF01261.27. The multiSMASH workflow^57^ was used to scan the downloaded Genbank files for *asbABCDE* homologs and tabulate the results. BGC presence and absence was mapped to the phylogenetic tree with iTOL v6.8.^30^ Default parameters were used for all software unless otherwise specified.^26^ A BGC region met the “AsbABCDE” rule if it contained matches to existing antiSMASH models for IucA_IucC (*asbA/asbB*), AMP-binding (*asbC*), and PP-binding (*asbD*), as well as the Pfam model for DUF6005 (*asbE*, PF19468.2).^56^ No *asbF* model was used due to a low bitscore between *mllF* and the expected Pfam model PF01261.27. The multiSMASH workflow^57^ was used to scan the downloaded Genbank files for *asbABCDE* homologs and tabulate the results. BGC presence and absence was mapped to the phylogenetic tree with iTOL v6.8.^30^ Default parameters were used for all software unless otherwise specified.

### DNA manipulation

Primers used for plasmid and strain construction are listed in Extended Data Figure 6. All fragments were amplified with Phusion DNA polymerase (New England Biolabs). Fragments for pAZ1 were fused using *in vivo* yeast DNA assembly.^58^ Competent *S. cerevisiae* HZ848 were freshly prepared and transformed with a mixture of purified PCR products and spread on uracil dropout medium (MilliporeSigma). Prototrophic colonies were grown in uracil dropout medium (MilliporeSigma) at 30 °C and 250 rpm and the plasmid was purified using the Yeast Plasmid MiniPrep II Kit (Zymo Research). The plasmid was transformed into TransforMax EPI300TM *E. coli* (Lucigen) and induced according to the manufacturer’s protocol. pAZ1 was purified using the BAC DNA Miniprep Kit (Zymo Research). Full plasmid sequencing (Primordium) confirmed the sequence of pAZ1 (Extended Data Fig. 6). pAZ1 was then electroporated into Δ*mxaF* to generate Δ*mxaF*/pAZ1. Fragments for *mll* deletion vector pMS2 were fused using Gibson Assembly Master Mix (New England Biolabs), transformed into competent DH5α *E. coli* (New England Biolabs), and purified using the GeneJET Plasmid Miniprep Kit (Thermo Fischer Scientific). Counterselection was achieved through patching onto Hypho succinate sucrose and Hypho kanamycin plates and selecting colonies with a successful double crossover event (growth on sucrose, no growth on kanamycin). The deletion of the *mll* cluster and absence of suppressor mutations was confirmed by whole genome sequencing (SeqCenter).

### Analytical scale identification of methylolanthanin from crude extracts

50 mL ½ Hypho media supplemented with 50 mM methanol and 1 μM Nd_2_O_3_ (99.99%, Chempur) was used to culture Δ*mxaF* and Δ*mxaF*/pAZ1 in 250 mL Erlenmeyer flasks to late-exponential phase (OD_600_ = 0.8). Supernatants were harvested by centrifugation at 4000xg at 4 °C. 25 mL of lyophilized supernatant equivalent were dissolved into 6 mL water and extracted using solid phase on HLB column preparation (MN Chromabond; 60 μM, 500 mg). HLB columns were conditioned with methanol (2 × 6 mL), then equilibrated with water (2 × 6 mL). Sample was loaded onto the column, the column was washed with water (3 × 6 mL), dried with vacuum and then eluted with methanol (2 × 6 mL). Methanol was removed *in vacuo*, the concentrated samples were dissolved in water/acetonitrile (50/50, 1 mL), and the sample was syringe filtered (0.2 μm PTFE).

A 2.5 μL sample was injected into an Agilent QTOF 6530 C coupled to an Agilent HPLC 1260 Infinity II instrument (G7115A 1260 DAD WR, G7116A 1260 MCI, G7167A 1260 multisampler, G7104C 1260 flexible pump). A C18 porous core column (Agilent Poroshell 120 EC-C18, 3.0 × 150 mm, 2.7 μm) was used for chromatography at 30ºC. The mobile phase consisted of solvent A (water + 0.1% formic acid (FA)) and solvent B (acetonitrile + 0.1% FA). The flow rate was set to 0.7 mL/min. After injection, the samples were eluted with the following method: 0–2 min 2% B, 2–20 min 2–98% B, followed by a 6-min washout phase at 98% B and a 3-min re-equilibration phase at 2% B. Data-dependent acquisition (DDA) of MS/MS spectra was performed in both positive and negative modes. ESI, MS, and MS/MS parameters are described in the Supplemental Methods.

### Feature finding in MZmine

MS/MS spectra were converted to .mzML files using MSConvert (*ProteoWizard*).^59^ MS1 feature extraction and MS/MS pairing were performed with MZmine 3.^33^ For positive mode data, an intensity threshold of 2.5E2 for MS1 spectra and of 0 for MS/MS spectra was used. MS1 ADAP chromatogram building was performed within a 20ppm mass window and a minimum highest intensity of 7.5E3 was set. XICs were deconvoluted using the local minimum search algorithm with a chromatographic threshold of 85%, a search minimum in RT range of 0.1 min, and a min ratio of peak top/edge of 1.8. Isotope features were grouped and features from different samples were aligned with 3 ppm mass tolerance and 0.05 min retention time tolerance. MS1 peak lists were joined using an m/z tolerance of 8 ppm and retention time tolerance of 0.4 min; alignment was performed by placing a weight of 75 on m/z and 25 on retention time. Gap filling was performed using an intensity tolerance of 20%, an m/z tolerance of 20 ppm, and a retention tolerance of 0.07 min. Feature areas and the corresponding MS/MS consensus spectra were exported as .csv and .mgf files respectively for post-processing using R and GNPS.

For negative mode data, an intensity threshold of 6E2 for MS1 spectra and of 0 for MS/MS spectra was used. MS1 ADAP chromatogram building was performed within a 20ppm mass window and a minimum highest intensity of 1.0E3 was set. XICs were deconvoluted using the local minimum search algorithm with a chromatographic threshold of 87.5%, a search minimum in RT range of 0.1 min, and a min ratio of peak top/edge of 1.8. Isotope features were grouped and features from different samples were aligned with 3 ppm mass tolerance and 0.08 min retention time tolerance. MS1 peak lists were joined using an m/z tolerance of 8 ppm and retention time tolerance of 0.4 min; alignment was performed by placing a weight of 75 on m/z and 25 on retention time. Gap filling was performed using an intensity tolerance of 20%, an m/z tolerance of 20 ppm, and a retention tolerance of 0.08 min. Feature areas and the corresponding MS/MS consensus spectra were exported as .csv and .mgf files respectively for post-processing using R Stuido^60^ and GNPS.^32^

### Classical molecular networking

All .mzML files were uploaded to the classical molecular networking workflow in GNPS for spectral networking and spectral library matching (gnps.ucsd.edu). For spectral library matching and spectral networking, the minimum cosine score to define spectral similarity was set to 0.7. The Precursor and Fragment Ion Mass Tolerances were set to 0.02 Da and Minimum Matched Fragment Ions to 4, Minimum Cluster Size to 1 (MS Cluster off). When Analog Search was performed the maximum mass difference was set to 100 Da. The GNPS job for positive analysis can be accessed through the following link: https://gnps.ucsd.edu/ProteoSAFe/status.jsp?task=c9fd99a7d4e147e5a281c23701ac0d2b (molecular family component index 753 / cluster index 182991) ; the negative analysis can be accessed through the following link: https://gnps.ucsd.edu/ProteoSAFe/status.jsp?task=d907c098d05a44b29ed7f670669c2a34 (molecular family component index 131 / cluster index 429623).

### Large scale cultivation, extraction, and purification of methylolanthanin

3 mL cultures of *ΔmxaF*/pAZ1 were grown in MP medium supplemented with 15 mM succinate and 50 μg/mL kanamycin overnight at 29 °C and 200 rpm in 14 mL vented cap culture tubes (Greiner). Multiple cultures were inoculated into 250 mL of MP medium supplemented with 15 mM succinate and 50 μg/mL kanamycin within a 500 mL Ultra Yield flask (Thompson Instrument) to a starting OD of 0.1 and were grown overnight at 29 °C and 200 rpm. The cells were separated by centrifugation (7 min, 1800xg, r.t.) and washed with ½ Hypho medium^45^ supplemented with 125 mM methanol and 50 μg/mL kanamycin. The cells were then resuspended in 5 mL of the same medium and up to five 1 L cultures (½ Hypho, 50 μg/mL kanamycin, 125 mM methanol, 1 μM Nd_2_O_3_) were inoculated at a starting OD of 0.1 in 2.5 L Ultra Yield flasks (Thompson Instrument). The cells were grown at 29 °C and 200 rpm until late stationary phase. The supernatant was obtained as a yellowish liquid after centrifugation (7 min, 15970xg, r.t.) and filtration (0.2 μm, PES).

For large scale solid phase extraction a *puriFlash®* plastic column (60 × 205 mm) was packed with HLB material (MN, 60 μm, 42 g). The column was equilibrated with methanol (2 × 500 mL) and water (2 × 500 mL). Then up to 2 L of supernatant was loaded onto the column and the column was washed with water (3 × 500 mL) and fully dried under nitrogen. The column was eluted with methanol (2 × 500 mL) into a 1 L round bottom flask. The solvent was removed *in vacuo* and the resulting residue was redissolved in water and lyophilised. By this method an average of 0.07 mg organic matter/mL supernatant was obtained.

HPLC purification was performed on an Agilent 1260 Infinity II preparative HPLC system (G7161A, G7165A, G1364E) equipped with an *Dr. Maisch* ReproSil Gold 120 C18 column (250 × 20 mm, 5 μm). The used solvent system was water/acetonitrile + 0.1% FA. The flow was set to 13.2 mL/min and the method was as follows: isocratic for 2 min at 98/2 water/acetonitrile, then in 40.8 min to 48/52 water/acetonitrile and in 5 min to 2/98 which was held for 10 min; retention time of methylolanthanin: 26 min (Extended Data Fig. 7). The concentration of a MLL stock solution was determined by qNMR which was then used to obtain the extinction coefficient *via* UV-vis (Supplementary Methods, Extended Data Fig. 8).

### NMR spectroscopy

NMR spectra were either recorded on a 800 MHz *Bruker* Avance III HD spectrometer equipped with a triple channel cryogenic probe at 298.15 K or on a 500 MHz *Bruker* BioSpin spectrometer equipped with a broad band observe 5-mm BB-H& CryProbe™ prodigy at 298.15 K. Spectra were recorded in 3 mm NMR tubes in a 9:1 H_2_O/ D_2_O (D_2_O 99.9%+0.03% TMSP, Deutero) mixture as well as, after lyophilisation and re-dissolving, in D_2_O only. If water suppression was applied the sequence zgesgp was used. All signals were assigned (Extended Data Fig. 5) using ^1^H, ^1^H-COSY, ^1^H, ^1^H-NOESY, ^1^H, ^1^H-TOCSY and phase-sensitive ^1^H, ^13^C-HSQC and ^1^H, ^13^C-HMBC experiments (Extended Data Fig. 9-10). Chemical shifts (*δ*) are reported in parts per million (ppm) using TMSP (0 ppm) as reference.

### Metal-binding experiments *via* mass spectrometry

Metal-binding experiments were performed using an Agilent QTOF 6530 C coupled to an Agilent HPLC 1260 Infinity II instrument (G7115A 1260 DAD WR, G7116A 1260 MCI, G7167A 1260 multisampler, G7104C 1260 flexible pump) *via* direct injection without a column and instead using a capillary (solvent system 1:1 water/acetonitrile + 0.1 FA, 0.7 mL/min flow) in positive mode. Samples were prepared by mixing methylolanthanin with an excess of LnCl_3_ (Ln = La, Nd, or Lu; purity: 99.99%) in a 1:1 methanol/water mixture.

### Elemental analysis of culture pellets

50 mL ½ Hypho media supplemented with 50 mM methanol and 1 μM Nd_2_O_3_ (99.99%, Chempur) was used to culture Δ*mxaF*, Δ*mxaF*/pAZ1, and Δ*mxaF*Δ*mll* in 250 mL Erlenmeyer flasks to late-exponential phase (OD600 = 0.8). Cells were pelleted at 4000xg and pellets were washed with water (3 × 10 mL) and resuspended in 1 mL of water and 1 mL of TraceMetal Grade 70% HNO_3_ (Thermo Fisher Scientific). Acidified cell pellets were boiled at 90 °C for 1 hr and pelleted at 4000xg. 1 mL of each sample was diluted with 19 mL of water. Samples were analyzed *via* ICP-MS (Laboratory for Environmental Analysis).

## DATA AVAILABILITY

The sequencing data from this study have been submitted to the NCBI Gene Expression Omnibus under the accession GSE244060. All MS .raw and centroided .mzML files are publicly available in the Mass spectrometry Interactive Virtual Environment (MassIVE) under massive.ucsd.edu with project identifiers MSV000090804 and MSV000083729, available at the following links: ftp://MSV000090804@massive.ucsd.edu and ftp://MSV000090284@massive.ucsd.edu Classical and feature based molecular networking for positive mode data can be accessed through gnps.ucsd.edu under the following direct links: https://gnps.ucsd.edu/ProteoSAFe/status.jsp?task=c9fd99a7d4e147e5a281c23701ac0d2b (classical MN) and https://gnps.ucsd.edu/ProteoSAFe/status.jsp?task=def58641630e488985bfbe5a15c94e32 (feature based MN). Classical and feature based molecular networking for negative mode data can be accessed through gnps.ucsd.edu under the following direct links: https://gnps.ucsd.edu/ProteoSAFe/status.jsp?task=d907c098d05a44b29ed7f670669c2a34 (classical MN) and https://gnps.ucsd.edu/ProteoSAFe/status.jsp?task=0478820eab694254b9eba0bbe4de2be6 (feature based MN).

All R code can be accessed at the following link: https://github.com/allegraaron/Lanthanophore_2023.

## ACKNOWLEDGEMENTS

The information, data, or work presented herein was funded in part by the Advanced Research Projects Agency-Energy, U.S. Department of Energy, under Award Number DE-AR0001337. The views and opinions of authors expressed herein do not necessarily state or reflect those of the United States Government or any agency thereof. This material is also based upon work supported by the National Science Foundation under Grants 2127732 (N.C.M.G.) and the ERC Starting grant Lanthanophore (L.J.D.). S.M.G.T. thanks Lisa Jobst, Jost Goormann and Alina Lobe for their contributions during their internships and the Studienstiftung des Deutschen Volkes for funding. A.T.A. was supported by the University of Denver (startup). A.M.Z. was supported by the U.S. Department of Defense (NDSEG).

## AUTHOR CONTRIBUTIONS

A.M.Z., S.M.G.T, L.J.D., and N.C.M.G. designed study. A.M.Z. generated, cultivated, measured growth of, and harvested RNA from genetic mutants, analyzed RNAseq data, prepared supernatants for mass spectrometry, and wrote paper. N.M.G. performed initial RNA-seq experiment identifying *mll*. D.P. provided access to and developed HRMS methods. A.T.A. and S.M.G.T performed mass spectrometry. A.T.A, S.M.G.T, L.J.D, and Z.L.R. elucidated structure of methylolanthanin. A.T.A. performed molecular networking and statistical analysis. S.M.G.T performed large scale cultivation, isolated MLL, determined MLL extinction coefficient, demonstrated metal binding properties, and designed, conducted, and analyzed NMR studies. M.T.P synthesized rhodopetrobactin B and methylolanthanin precursors as references for verification. Z.L.R. performed bioinformatic analyses. A.M.Z., S.M.G.T, A.T.A, and Z.L.R contributed text and figures. L.J.D. and N.C.M.G acquired funding and access for research, infrastructure, provided logistical support, and edited paper

