## Supplemental text and table for "Discovery and characterization of the first known biological lanthanide chelator"

### SUPPLEMENTARY INFORMATION

#### SUPPLEMENTARY METHODS

##### Identification of methylolanthanin at analytical scale from crude extracts

ESI parameters were set to the following: (positive mode) Gas Temp: 250 °C, Drying Gas: 11 L/min, Nebuliser 45 psi, Sheath Gas Temp 350 °C, Sheath Gas Flow 12 L/min; VCap 3500 V, Fragmentor 100 V, Skimmer 65 V, Oct 1 RF Vpp 750 V; mass range: 100-1700; rate 5 spectra/s; time 200 ms/spectrum; transients/spectrum 2641; Reference masses used: 121.0509 and 922.0098; (negative mode) Gas Temp: 275 °C, Drying Gas: 11 L/min, Nebuliser 35 psi, Sheath Gas Temp 350 °C, Sheath Gas Flow 12 L/min; VCap 3000 V, Fragmentor 100 V, Skimmer 65 V, Oct 1 RF Vpp 750 V; rate 5 spectra/s; time 200 ms/spectrum; transients/spectrum 2677; Reference masses used: 112.9855 and 1033.9881.

MS/MS settings for iterative aMS/MS were the following and were measured from 2 to 22 min: Spectral Parameters: (MS 100-1700 m/z; rate 4 spectra/s; time 250; transients/spectrum 3330; MS/MS 50-1000 m/z, rate 4, time 250, transients/spectrum 3211; isolation width ~1.2 m/z); cycle time 0.85 seconds

Collision Energy - fixed: 10, 20 and 40 V; Precursor Selection I : 2 per cycle; Threshold 2000 counts (relative Threshold 0.01%); Active Exclusion after 2 Spectra, released after 0.1 min; (the reference molecules were excluded); Settings for iterative MS/MS: mass tolerance +/- 20 ppm; retention time exclusion tolerance +/-0.2 min; Precursor Selection II: Isotope Model: unbiased; Charge State selection: 1, 2, Unk; Abundance Dependent Accumulation - Scan Speed varied based on precursor abundance.

##### Solid phase extraction for Q Exactive HF

50 mL of lyophilized supernatant were reconstituted in 6 mL of 3% methanol/LCMS grade water and was extracted onto HLB cartridges (MN Chromabond; 60 µM, 500 mg). HLB cartridges were pre-activated with methanol (2x3 mL), then were washed with water + 3% methanol (2x3 mL). Samples were loaded dropwise onto SPE cartridges before cartridges were washed with water+3%methanol (2x3 mL). Then samples were eluted stepwise into 2 mL of 50% methanol, 2 mL of 80% methanol, 2 mL of 100% methanol. Samples were weighed and reconstituted with 100 µL 80% methanol/20% water.

##### UHPLC-MS/MS on Q Exactive HF

For data-dependent UHPLC-MS/MS analysis, 5 µL of sample were injected per run. A C18 EVO porous core column (Kinetex C18, 50 × 2 mm, particle size of 1.8 µm, pore size of 100 Å, Phenomenex) was used for reversed-phase chromatography. A Vanquish high-pressure binary gradient system couple to a Q Exactive HF mass spectrometer and method according to Stincone *et al.* was used. The mobile phase consisted of solvent A (water+ 0.1% formic acid (FA)) and solvent B (acetonitrile (ACN) + 0.1% FA), and the flow rate was 0.5 ml $min^{-1}$. After injection, the samples were eluted with the linear gradient: 0–8 min 5–50% B, followed an increase from 50-99 % B from 8-12 min, and by a 3-min washout phase at 99% B and a 3-min re-equilibration phase at 5% B. Data-dependent acquisition (DDA) of MS/MS spectra was performed in positive mode. ESI parameters were set to a sheath gas flow of 50AU, auxiliary gas flow of 12 AU, sweep gas flow of 1 AU and auxiliary gas temperature of 400 °C, while the spray voltage was set to 3.5 kV, the inlet capillary to 250 °C. The MS scan range was set to 150–1,500 m/z with a resolution at m/z 200 (Rm/z 200) of 120,000. The maximum ion injection time was set to 100 ms with an automated gain control (AGC) target of $1.0 \times 10^6$. Up to five MS/MS spectra per MS1 survey scan were recorded in DDA mode with Rm/z 200 of 15,000 with one micro-scan. The maximum ion injection time for MS/MS scans was set to 50 ms with an AGC target of $5 \times 10^5$ ions. The MS/MS precursor isolation window was set to m/z 1. The normalized collision energy was set to a stepwise increase from 25 to 35 to 45% with $z = 1$ as default charge state. MS/MS scans were triggered at the apex of chromatographic peaks within 2 to 15 s from their first occurrence. Dynamic precursor exclusion was set to 5 s. Ions with unassigned charge states were excluded from MS/MS acquisition as well as isotope peaks.

**Feature based molecular networking**

The .csv and .mgf file outputs from Feature finding in MZmine were directly uploaded to the feature based molecular networking workflow in GNPS for spectral networking and spectral library matching (gnps.ucsd.edu). For spectral library matching and spectral networking, the minimum cosine score to define spectral similarity was set to 0.7. The Precursor and Fragment Ion Mass Tolerances were set to 0.02 Da and Minimum Matched Fragment Ions to 4, Minimum Cluster Size to 1 (MS Cluster off). When Analog Search was performed the maximum mass difference was set to 100 Da. The GNPS job for positive analysis can be accessed through the following link: https://gnps.ucsd.edu/ProteoSAFe/status.jsp?task=def58641630e488985bfbe5a15c94e32; the negative analysis can be accessed through the following link: https://gnps.ucsd.edu/ProteoSAFe/status.jsp?task=0478820eab694254b9eba0bbe4de2be6.

**LC-MS/MS data cleanup and statistics**

The .csv feature tables from Feature finding in MZmine 3 were blank subtracted, imputed, normalized, and scaled using R code available at https://github.com/allegra-aron/Lanthanophore_2023/. A media blank was used for blank subtraction, specifically features with mean intensity of 30% abundance in the media as compared to samples were removed. Imputation was utilized to remove zero values by replacing them with the limit of detection for the experiment. Normalization was performed by scaling the intensities of all features in a sample by the total ion current (or sum) of the sample (total ion current or TIC normalization). Volcano plots and Kruskal-Wallis analysis followed by pairwise Wilcoxon tests and Benjamini-Hochberg (BH) correction were performed using R code provided in the same location (https://github.com/allegra-aron/Lanthanophore_2023/). The blank subtracted, imputed, TIC normalized table was utilized for volcano plots and Kruskal-Wallis analysis. Principal component analysis was performed using the scaled data.

**Concentration determination *via* qNMR and determination of the extinction coefficient**

$^1$H qNMR was measured in 3 mm NMR tubes (*Deutero*, D800-3-7) on a *Bruker* Avance III HD spectrometer equipped with a triple channel cryogenic probe operating at the Larmor frequency of

800 MHz (18.8 T) at 298.15 K. The following acquisition settings were used: spectral width of 12820.5 Hz, 48 scans, acquisition time of 1.278 s, 60 s relaxation delay and 16 k real points. For water suppression the pulse sequence zgesgp was used. A stock solution (10 mg/mL; MW: 130.11 g/mol) in $D_2O$ (+ 0.03% TMSP, *Deutero*) of the quantitative reference material calcium formate (*Merck*; certified reference material, 99.58%) was gravimetrically prepared (*Mettler Toledo* UMT2 precision scale, Δ=0.1µg) and diluted 1:10 with $D_2O$ + 0.03% TMS before it was mixed 1:10 with an aqueous solution with an unknown concentration of methylolanthanin. The equation (1) was used to calculate the concentration ($c_x$) of methylolanthanin. With *I* standing for integral, *N* for number of atomic nuclei and $P_{Std}$ for purity (%) . The qNMR spectrum and the used integral values are shown in Extended Data Fig. 8.

$$(1)\ c_x = \frac{I_x}{I_{Std}} \times \frac{N_{Std}}{N_x} \times c_{Std} \times P_{Std}$$

With this the concentration of the aqueous solution of methylolanthanin was determined and was usedto prepare a series of samples in water with known concentration to determine the extinction coefficient (ε) using the *Beer-Lambert* law (Extended Data Fig. 8). UV-vis spectra were recorded on an *Agilent* Cary 60 spectrometer at 25 °C in *Brand* micro cuvettes (10 mm pathlength; 70-500 µL) with a scan rate of 600 nm/min in 1 nm steps. All spectra were measured using the automatic baseline correction after measuring the used solvent. In order to determine the extinction coefficient, the absorbance at 251 nm was plotted against the concentration which gave an extinction coefficient of 25 +/– 0.4 $mM^{-1}cm^{-1}$.

#### SUPPLEMENTARY TABLES

**Supplementary Table 1 | Prevalence of *mll* in *Methylobacteriaceae***

| #Assembly | Strain | AsbABCDE present? |
|---|---|---|
| GCF_001741865.1 | Bosea vaviloviae Vaf18 | False |
| GCF_004802635.2 | Methylocystis heyeri H2 | False |
| GCF_009363855.1 | Microvirga thermotolerans HR1 | False |
| GCF_016804325.1 | Methylobacterium aquaticum BG2 | True |
| GCF_001548015.1 | Methylobacterium aquaticum MA-22A | False |
| GCF_021228015.1 | Methylobacterium currus TP-3 | True |
| GCF_003173715.1 | Methylobacterium durans 17SD2-17 | False |
| GCF_029761955.1 | Methylobacterium indicum JDJ13 | False |
| GCF_017347565.1 | Methylobacterium indicum VL1 | False |

| | | |
|---|---|---|
| GCF_000364445.2 | Methylobacterium mesophilicum SR1.6/6 | False |
| GCF_029714205.1 | Methylobacterium nodulans CB376 | False |
| GCF_000022085.1 | Methylobacterium nodulans ORS 2060 | False |
| GCF_022533465.1 | Methylobacterium organophilum WPA_B | False |
| GCF_000757795.1 | Methylobacterium oryzae CBMB20 | False |
| GCF_021398735.1 | Methylobacterium oryzae H33R-06 | False |
| GCF_001936175.1 | Methylobacterium phyllosphaerae CBMB27 | False |
| GCF_003173735.1 | Methylobacterium radiodurans 17Sr1-43 | False |
| GCF_000019725.1 | Methylobacterium radiotolerans JCM 2831 | False |
| GCF_021484845.1 | Methylobacterium radiotolerans NYY1 | False |
| GCF_003173775.1 | Methylobacterium sp. 17Sr1-1 | False |
| GCF_029691625.1 | Methylobacterium sp. 391_Methyba4 | False |
| GCF_000019365.1 | Methylobacterium sp. 4-46 | False |
| GCF_001542815.1 | Methylobacterium sp. AMS5 | True |
| GCF_001854385.1 | Methylobacterium sp. C1 | False |
| GCF_025813715.1 | Methylobacterium sp. FF17 | False |
| GCF_028583545.1 | Methylobacterium sp. NMS14P | False |
| GCF_008000895.1 | Methylobacterium sp. WL1 | False |
| GCF_003254375.1 | Methylobacterium sp. XJLW | False |
| GCF_023546765.1 | Methylobacterium tardum DSM 19566 | False |
| GCF_003173755.1 | Methylobacterium terrae 17Sr1-28 | False |

| | | | |
|---|---|---|---|
| GCF_022179725.1 | Methylorubrum aminovorans NBRC 15686 | True | |
| GCF_000022685.1 | Methylorubrum extorquens AM1 | True | |
| GCF_030062705.1 | Methylorubrum extorquens ATCC 55366 | True | |
| GCF_000021845.1 | Methylorubrum extorquens CM4 | True | |
| GCF_000083545.1 | Methylorubrum extorquens DM4 | True | |
| GCF_026122615.1 | Methylorubrum extorquens NBC_00036 | True | |
| GCF_026122595.1 | Methylorubrum extorquens NBC_00404 | True | |
| GCF_030255395.1 | Methylorubrum extorquens PA1 | True | |
| GCF_000018845.1 | Methylorubrum extorquens PA1 | True | |
| GCF_001971665.1 | Methylorubrum extorquens PSBB040 | True | |
| GCF_900234795.1 | Methylorubrum extorquens TK 0001 | ** | Fragmented AsbC |
| GCF_022179745.1 | Methylorubrum podarium DSM 15083 | True | |
| GCF_000019945.1 | Methylorubrum populi BJ001 | True | |
| GCF_002355515.1 | Methylorubrum populi P-1M | True | |
| GCF_006740745.1 | Methylorubrum populi YC-XJ1 | True | |
| GCF_014199985.1 | Methylorubrum rhodesianum DSM 5687 | True | |
| GCF_014199935.1 | Methylorubrum rhodinum DSM 2163 | False | |
| GCF_900114375.1 | Methylorubrum salsuginis CGMCC 1.6474 | False | |
| GCF_021117295.1 | Methylorubrum sp. B1-46 | True | |

| | | |
|---|---|---|
| GCF_024347855.1 | Methylorubrum sp. GM97 | True |
| GCF_022179765.1 | Methylorubrum suomiense DSM 14458 | False |
| GCF_022179785.1 | Methylorubrum thiocyanatum JCM 10893 | True |
| GCF_014845115.1 | Methylorubrum zatmanii LMG 6087 | False |
