## Supplemental Figures for "Discovery and characterization of the first known biological lanthanide chelator"

### EXTENDED DATA FIGURES AND TABLES

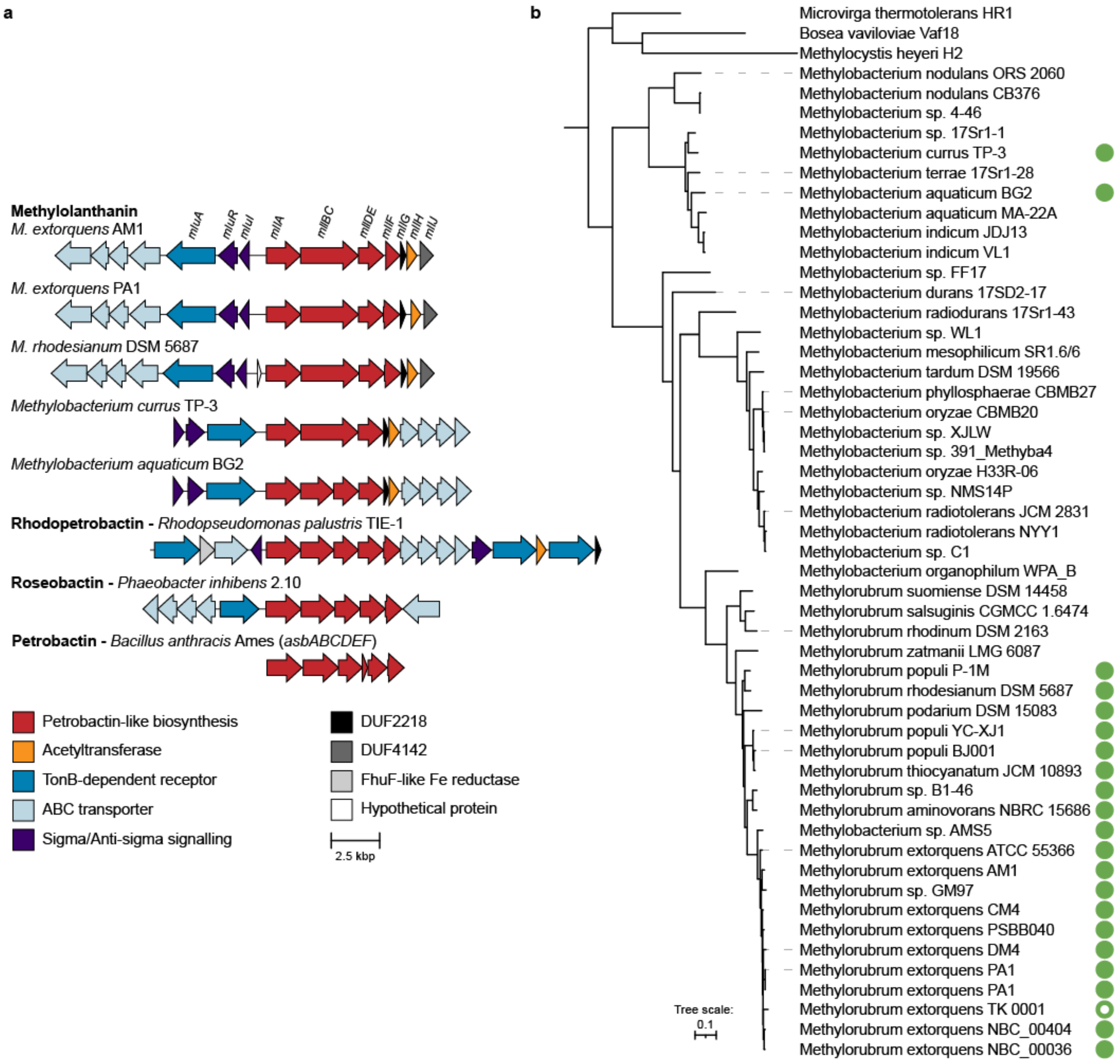


**Extended Data Fig. 1 | Comparative genomics of *mll* and prevalence in *Methylobacteriaceae*. a,** comparison of *mll* from *Methylorubrum exotorquens* AM1 against homologous clusters, including BGCs encoding the characterized siderophores rhodopetrobactin,[25] roseobactin,[59] and petrobactin.[26] Genes are drawn to scale. **b,** a maximum-likelihood phylogenetic species tree of *Methylorubrum* and *Methylobacterium* strains, annotated by the presence of a BGC homologous to *mll* (filled green circle). The *asbC* homolog in *M. extorquens* TK 0001 (open circle) did not match the AMP-binding domain (PF00501) and may be non-functional. Assembly accession numbers are provided in Supplementary Table 1.

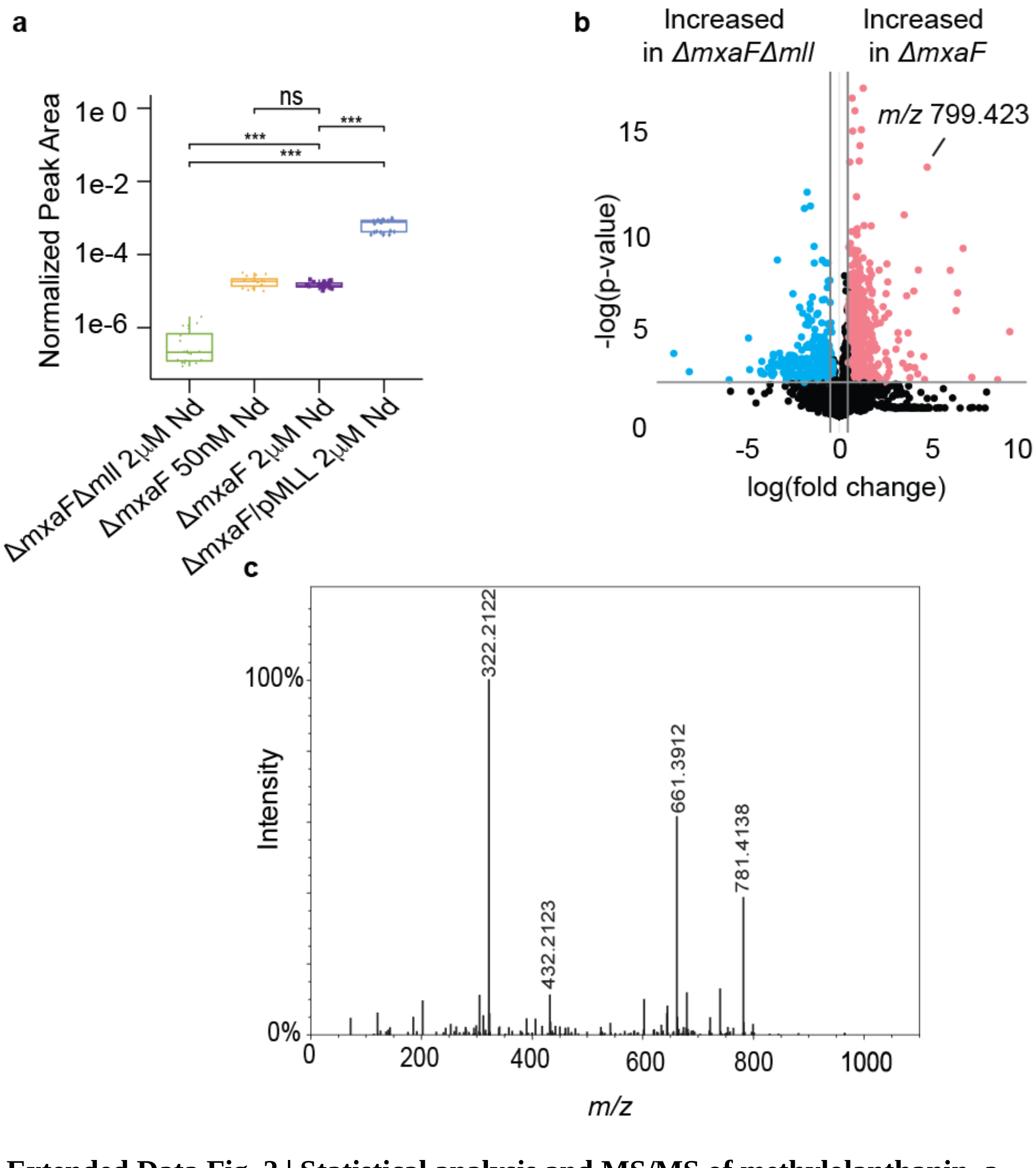


**Extended Data Fig. 2 | Statistical analysis and MS/MS of methylolanthanin. a**, Boxplots of the normalized peak area of feature *m/*z 799.4232 in positive mode reveal signifigant differences between Δ*mxaF* and Δ*mxaF*Δ*mll* and between Δ*mxaF*/pMLL but differences between Δ*mxaF* 2 µM Nd and Δ*mxaF* 50 nM were not significant. Statistics were calculated with n=5 per group. Kruskal-Wallis, followed by pairwise Wilcoxon tests and Benjamini-Hochberg (BH) correction, was used (***p < 0.001). **b**, *m/z* 799.4232 is one of the most significantly increased features in Δ*mxaF* versus Δ*mxaF*Δ*mll* supernatants when analyzed using ESI-UPLC-MS/MS in negative ionization mode. Volcano plot analysis was performed with n=20 per group. Fold changes 0.5 and a p-value < 0.01 are highlighted. **c**, The MS/MS spectrum in positive mode of the 1+ methylolanthanin.

**a**

| $^{1}H$ NMR chemical shifts (ppm) for Methylolanthanin | | | | |
|---|---|---|---|---|
| C/H | $\delta_H$ ($D_2O$) | Mult. (J in Hz) | $\delta_C$ ($D_2O$) | HMBC |
| 1 | - | - | 162.6 | |
| 2 | 6.94 | 8.6 (d) | 118.3 | C1, C2, C4 |
| 3 | 7.67 | 8.6 (d) | 132.3 | C1, C3, C5 |
| 4 | - | - | 128.7 | |
| 5 | - | - | 173.4 | |
| 6 | - | - | - | - |
| 7 | 3.36, 3.38 | m | 48.3, 42.2 | C5, C8, C9 |
| 8/9 | 1.47, 1.57; 1.42, 1.58 | m, m | 27.1; 28.6 | |
| 10 | 3.29, 3.33 | m | 51.8 | C11 |
| 11 | 3.25, 3.30 | m | 48.6 | |
| 12 | - | - | 176.5 | |
| 13 | 2.07, 2.07 | s,s | 23.3 | C12 |
| 14 | 1.48, 1.56 | m | 27.2, 27.3 | |
| 15 | 1.39, 1.43 | m | 28.7, 28.6 | |
| 16 | 3.08, 3.09 | m | 41.7 | C18 |
| 17 | - | - | - | |
| 18 | - | - | 175.1 | |
| 19 | $H_A$ 2.66<br>$H_B$ 2.56 | 14.4 (d, 4 AB systems overlapping) | 47.2 | C18, C19, C20, C21 |
| 20 | - | - | 77.9 | |
| 21 | - | - | 182.0 | |
| C/H | $\delta_H$ ($H_2O/D_2O$) | Mult. (J in Hz) | $\delta_C$($H_2O/D_2O$) | HMBC |
| 6 | 8.31, 8.28 | 6.3 (t), 6.2 (t) | - | |
| 17 | 7.81, 7.80 | 5.9 (t), 6.0, (t) | - | |
| 19 | $H_A$ 2.68<br>$H_B$ 2.57 | 14.7 (d, 4 AB systems overlapping) | 47.2 | C18, C19, C20, C21 |

**b**

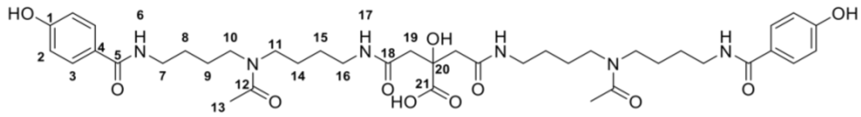

**Extended Data Fig. 3 | NMR assignments of methylolanthanin.** **a**,$^{1}H$ NMR chemical shifts (ppm) for methylolanthanin. **b**, Structure of methylolanthanin with numbering for NMR assignments.

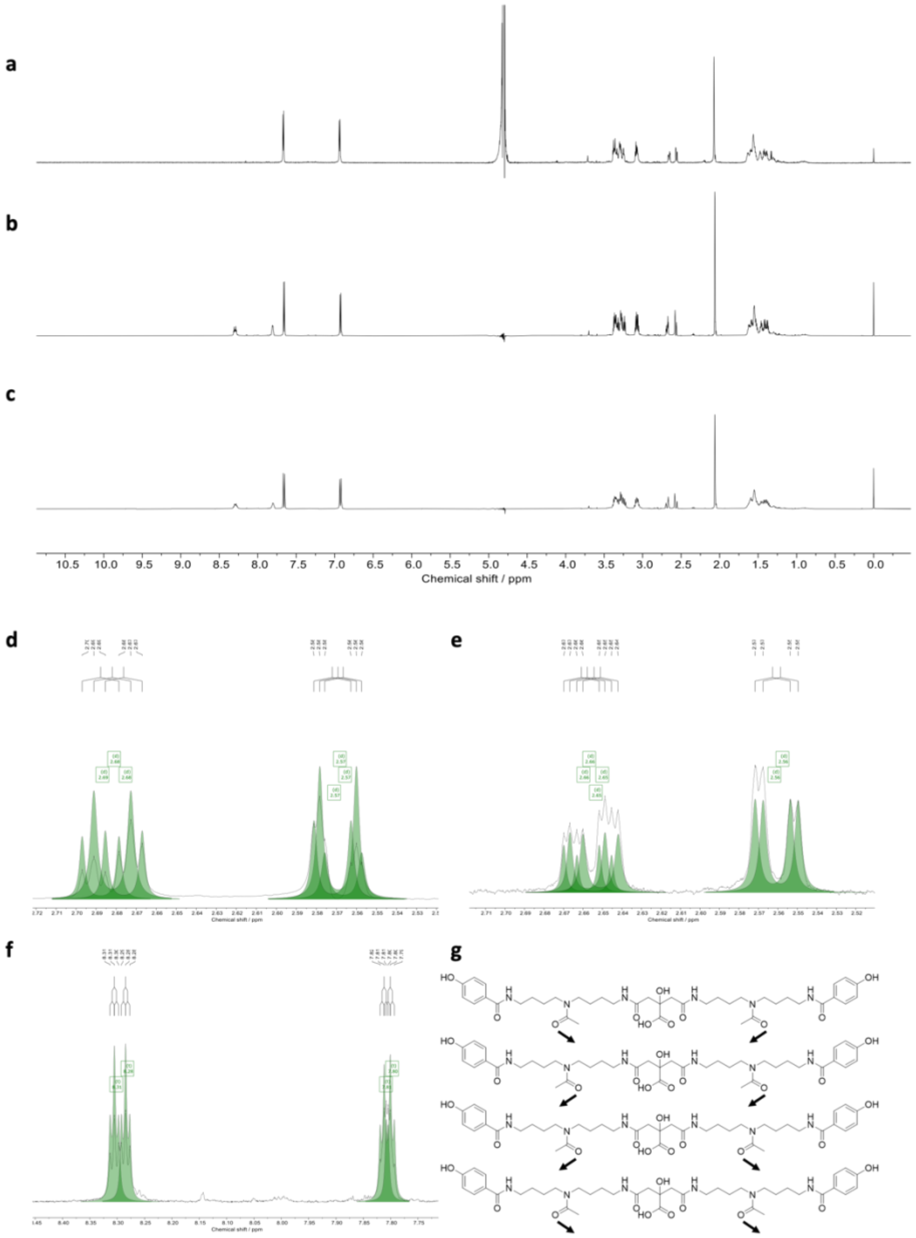


**Extended Data Fig. 4 | [1]H NMR spectra of methylolanthanin**. **a,** in $D_2O$ (800 MHz), **b,** in $H_2O/D_2O$ (9:1)(800 MHz) and **c,** in $H_2O/D_2O$ (9:1) (500 MHz). **d,** Deconvoluted signals observed for the four overlapping AB spin systems of the diastereotopic methylene signals of the central citric acid moiety of methylolanthanin in $D_2O/H_2O$ (800 MHz) and **e,** in $D_2O$ (800 MHz). **f,** deconvoluted signals observed for the secondary amines protons of methylolanthanin in $D_2O/H_2O$ (800 MHz) caused by the proximity to differently orientated acyl groups. **g,** Molecular structures of the four possible conformers of methylolanthanin causing the signal splitting.

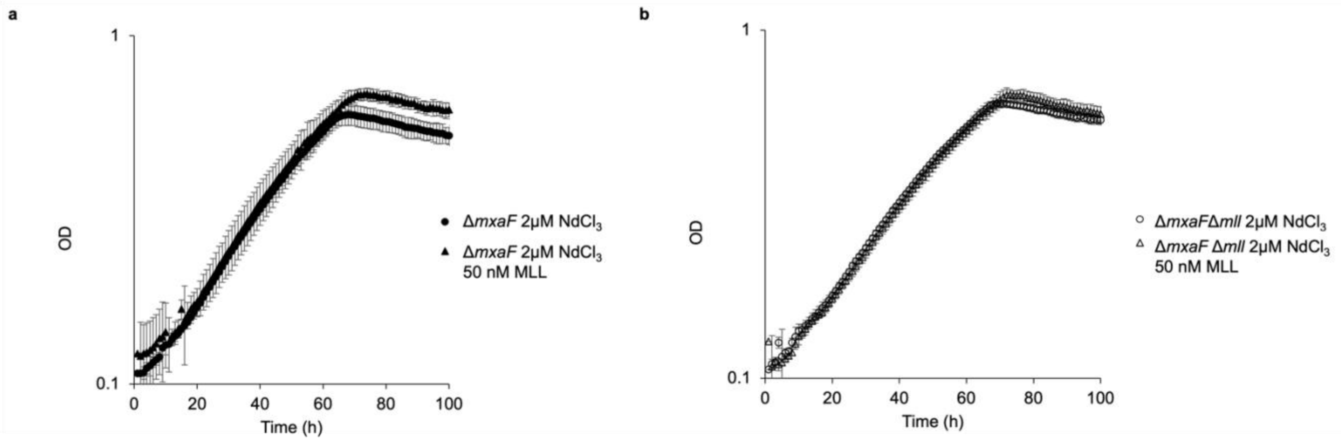


**Extended Data Fig. 5 | Exogenous methylolanthanin affects growth of Δ*mxaF* and Δ*mxaF*Δ*mll*.** **a**, Addition of methylolanthanin to Δ*mxaF* cultures (filled triangles) significantly increases growth yield ($p = 0.037$) compared to cultures grown without methylolanthanin (filled circles). Individual data points represent the mean of three replicates. **b**, Addition of methylolanthanin to Δ*mxaF*Δ*mll* cultures (open triangles) significantly increases growth yield ($p = 0.036$) compared to cultures grown without methylolanthanin (open circles). Individual data points represent the mean of three replicates.

a

| Primer | Sequence (5' to 3') | Plasmid | Product |
|---|---|---|---|
| AZ09 | ACGAGAAATCTTGCCGTTGGC | pAZ1 | META1p4132-META1p4133 + first 500 bp of META1p4134 |
| AZ10 | ATCAACCCGGATCTGCGCC | pAZ1 | last 500 bp of META1p4133 + META1p4134-META1p4138 |
| AZ11 | GAGAACGAGATGACGTTGGA | pAZ1 | joint fragment: 200 bp HZ848 backbone + 200 bp AM1 backbone |
| AZ12 | ATTGCCATACCTTTGGTCGTTTTTTAGCCGCTAAAACGG | pAZ1 | joint fragment: 200 bp HZ848 backbone + 200 bp AM1 backbone / HZ848 backbone |
| AZ13 | CGACCAAAGGTATGGCAATTTAGAAAAACTCATCGAGCATCA | pAZ1 | joint fragment: 200 bp HZ848 backbone + 200 bp AM1 backbone / AM1 backbone |
| AZ14 | TCCTTTTAACAGCGATCGCG | pAZ1 | joint fragment: 200 bp HZ848 backbone + 200 bp AM1 backbone |
| AZ15 | CATTCGATAAGCCACGAAATATGACCATGATTACGCCAAG | pAZ1 | joint fragment: 200 bp AM1 backbone + 200 bp EPI300 backbone |
| AZ16 | TTTACCAAGGCATCATTGCATGTAAGCGGATGCCGG | pAZ1 | joint fragment: 200 bp AM1 backbone + 200 bp EPI300 backbone / / AM1 backbone |
| AZ17 | TGCAATGATGCCTTGGTAAAATCTACGTCTGTCGAGAAGT | pAZ1 | joint fragment: 200 bp AM1 backbone + 200 bp EPI300 backbone / EPI300 backbone |
| AZ18 | TTGATATCGGGGGTTAGTTCG | pAZ1 | joint fragment: 200 bp AM1 backbone + 200 bp EPI300 backbone |
| AZ19 | GTGTTCACCCTTGTTACACCGT | pAZ1 | joint fragment: 200 bp EPI300 backbone + first 200 bp META1p4132 |
| AZ20 | TGAATTGGACTTAACGATTCAGGGCACCAATAACTGCCT | pAZ1 | joint fragment: 200 bp EPI300 backbone + first 200 bp META1p4132 / EPI300 backbone |
| AZ21 | GAATCGTTAAGTCCAATTCAGATCATGCCACCCTCGTCTC | pAZ1 | META1p4132-META1p4133 + first 500 bp of META1p4134 |
| AZ22 | ACGAGAAATCTTGCCGTTGGC | pAZ1 | joint fragment: 200 bp EPI300 backbone + first 200 bp META1p4132 |
| AZ23 | CTCGTTGGCGGTCTGGTCG | pAZ1 | joint fragment: last 200 bp META1p4138 + 200 bp HZ848 backbone |
| AZ24 | GTAGATTCGGCATTAGGCAACGGTTGCCACCTTTCGGGAC | pAZ1 | last 500 bp of META1p4133 + META1p4134-META1p4138 / joint fragment: last 200 bp META1p4138 + 200 bp HZ848 backbone |
| AZ25 | TTGCCTAATGCCGAATCTACGAGGCCAAAAAGCTCGCTTTCAG | pAZ1 | joint fragment: last 200 bp META1p4138 + 200 bp HZ848 backbone / HZ848 backbone |
| AZ26 | TGCCGGCACGTTAACCG | pAZ1 | joint fragment: last 200 bp META1p4138 + 200 bp HZ848 backbone |
| AZ27 | ATTTCGTGGCTTATCGAATGAACAAACCACCGCTGGTAGC | pAZ1 | AM1 backbone |
| MS25 | ACTTGGTCTGACAGTTACCATCTCCCCGCATATCGACTAC | pMS2 | *mll* upstream flank |
| MS26 | GGGTTTGCTGCCGGGATTCAATGGCCCGTATGGTCACGTT | pMS2 | *mll* downstream flank |
| MS29 | CTTAACATCCACGGCTCCTAATGTGCAGGTTGTCGGTGTC | pMS2 | pCM433kanT backbone |
| MS30 | GTAGTCGATATGCGGGGAGATGGTAACTGTCAGACCAAGT | pMS2 | pCM433kanT backbone |

b

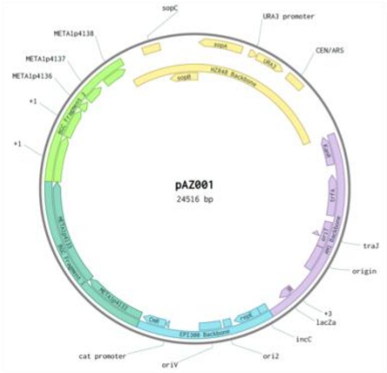


c

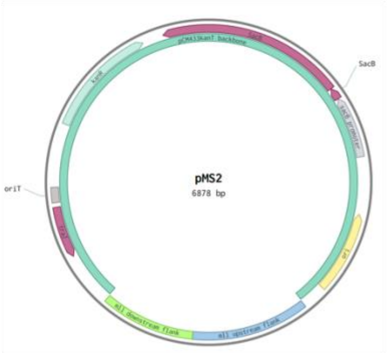


**Extended Data Fig. 6 | Primers and plasmids used in study. a**, Primers used to construct pAZ1 and pMS2. **b**, Plasmid map of pAZ1used to generate the *mll* overexpression mutant. **c**, Plasmid map of pMS2 used to generate the *mll* deletion mutant.

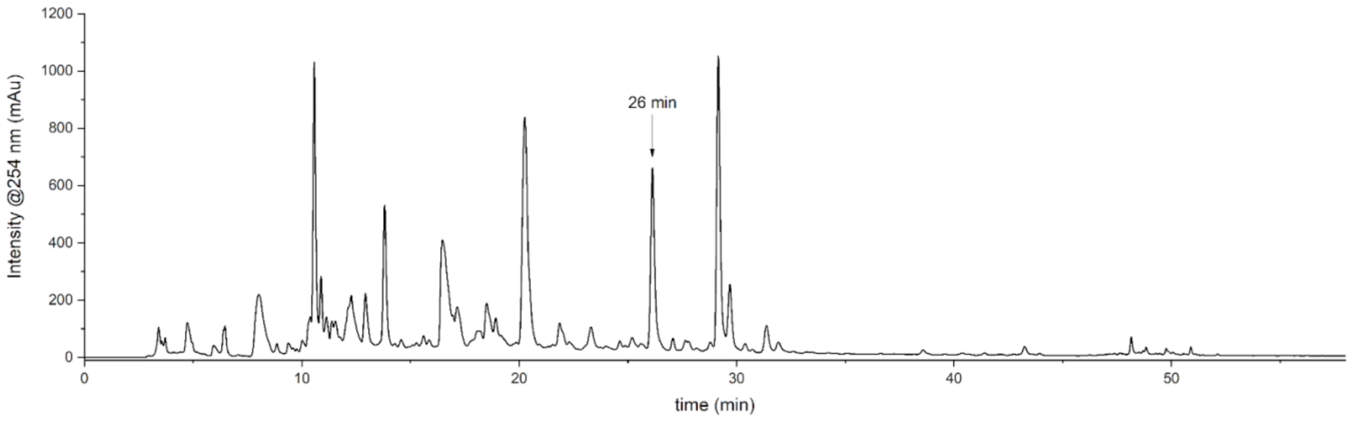


**Extended Data Fig. 7 | Preparative HPLC chromatogram of methylolanthanin.** Extraction from culture supernatants ran on reversed-phase column shows elution of methylolanthanin at 26 minutes.

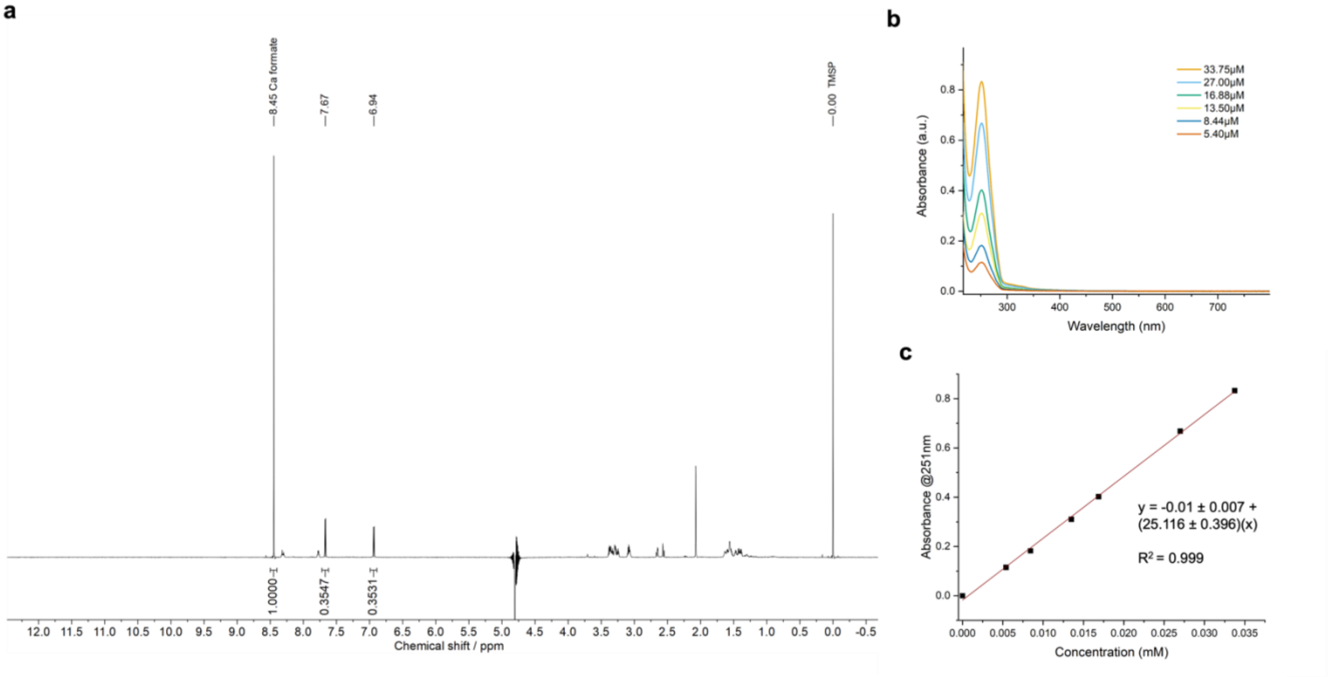


**Extended Data Fig. 8 | qNMR and extinction coefficient determination of methylolanthanin.** **a**, qNMR of methylolanthanin in $H_2O/D_2O$ (9:1) (800MHz). **b**, UV-vis spectra of methylolanthanin at different concentrations in water. **c**, Concentration of methylolanthanin plotted against the absorbance at 251 nm for determination of the extinction coefficient from regression line.

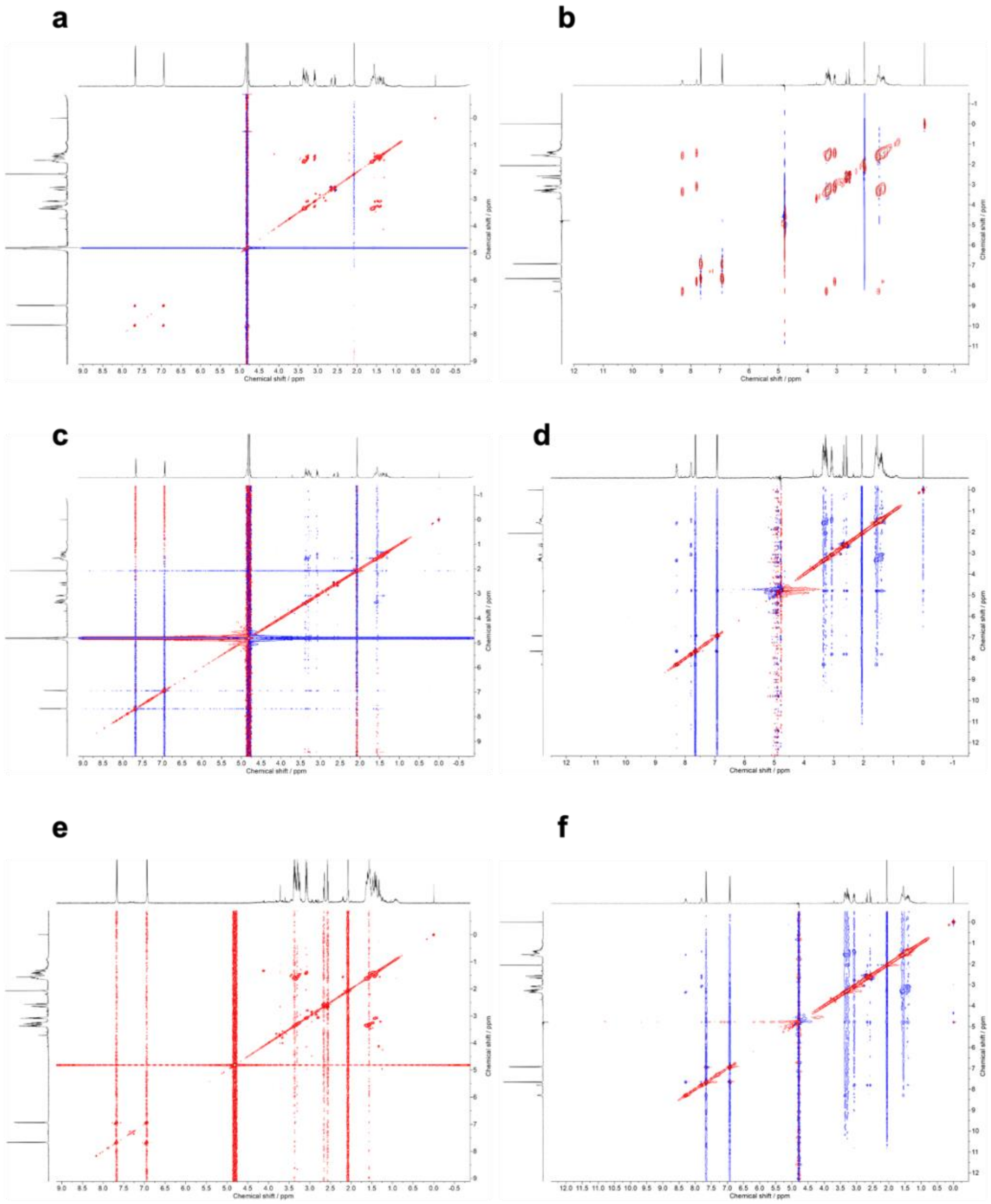


**Extended Data Fig. 9 | 2D proton NMR spectra of methylolanthanin**. **a,** TOCSY spectrum (800 MHz, $D_2O$). **b,** TOCSY spectrum (500 MHz, 9:1 $H_2O/D_2O$measured with water suppression. **c,** NOESY spectrum (800 MHz, $D_2O$). **d,** NOESY spectrum (500 MHz, 9:1 $H_2O/D_2O$) measured with water suppression. **e,** COSY spectrum (800 MHz, $D_2O$). **f**, ROESY (500 MHz, 9:1 $H_2O/D_2O$) spectrum measured with water suppression.

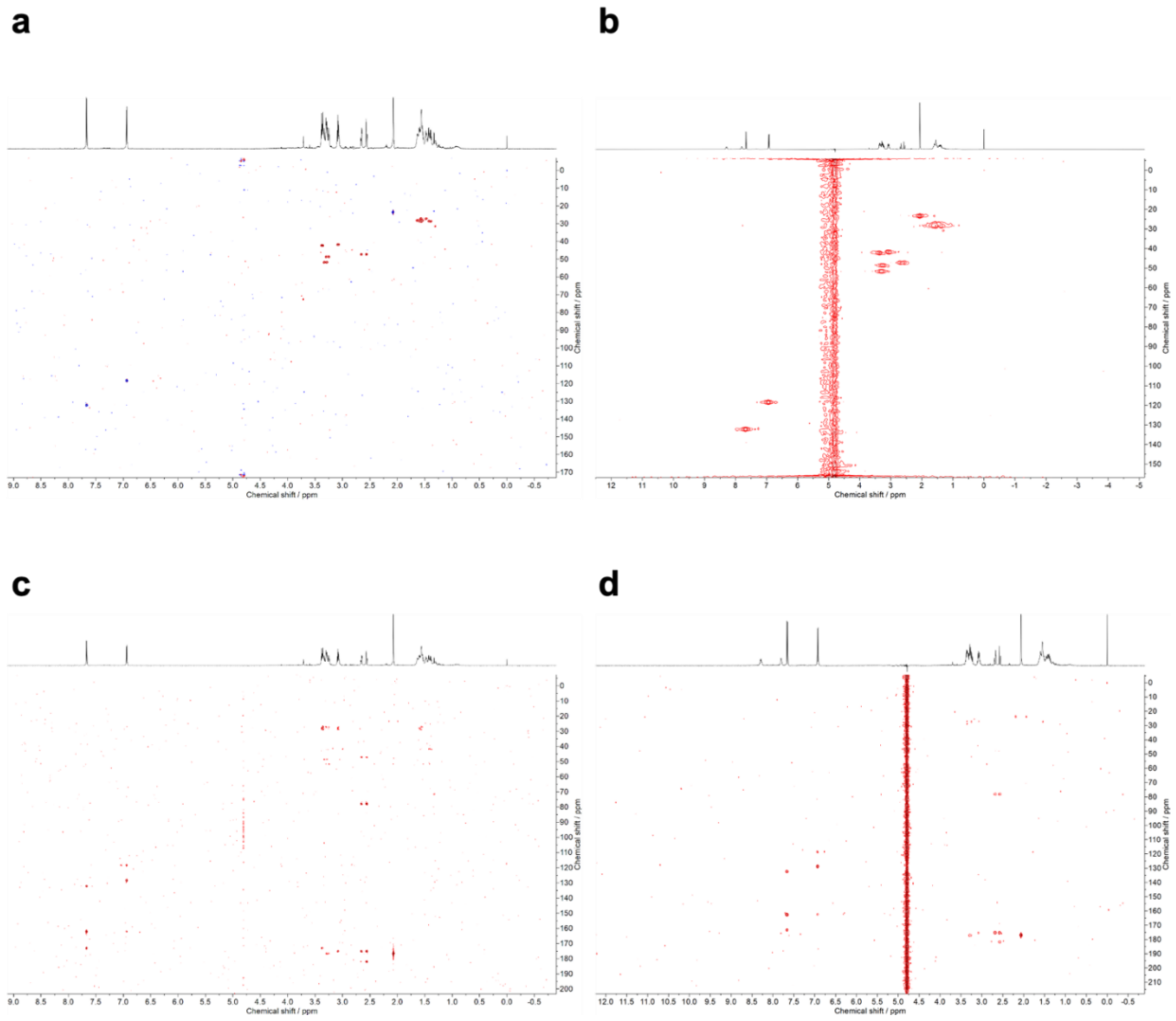


**Extended Data Fig. 10 | Heteronuclear $^{1}H$-$^{13}C$ 2D NMR spectra of methylolanthanin**. **a**, phase-sensitive HSQC (800 MHz, $D_2O$) spectrum **b**, HSQC (500 MHz, 9:1 $H_2O/D_2O$) spectrum measured with water suppression **c**, HMBC (800 MHz, $D_2O$). **d**, HMBC (500 MHz, 9:1 $H_2O/D_2O$) spectrum measured with water suppression.
